# RNA at Lipid/Water Interfaces: Molecular Insights from Coarse-Grained Simulations and Reflectivity Data Predictions

**DOI:** 10.1101/2025.11.21.689668

**Authors:** Mohd Ibrahim, Jürgen Köfinger, Martin Zacharias, Emanuel Schneck, Nadine Schwierz

## Abstract

Interactions between RNA and lipids are fundamental for biological processes and are increasingly exploited for RNA delivery by lipid nanoparticles. However, RNA-lipid interactions remain challenging to characterize at the molecular level. Here, we address the modeling of RNA at lipid/water interfaces using coarse-grained (CG) simulations, experimental validation using scattering data and prediction of neutron (NR) and X-ray reflectivity (XRR) profiles from the simulations. Using neutral DOPC and cationic DOTAP bilayers, we show that lipid-RNA interactions depend strongly on RNA secondary structure, with single-stranded regions exhibiting higher interfacial affinity than double-stranded segments. We validate the CG lipid simulations, showing that while they reproduce experimental X-ray scattering data only qualitatively, the agreement improves markedly after backmapping to atomistic resolution followed by energy minimization and short all-atom molecular dynamics simulations. We further simulated distinct tRNA conformations and analyzed the influence of RNA secondary structure, concentration, solvent contrast, and lipid deuteration on NR and XRR signals identifying the conditions under which such experiments probe RNA adsorption and discriminate between different RNA conformations. Together, these results demonstrate that CG simulations combined with reflectivity data provide a powerful approach to probe RNA adsorption and structure at lipid/water interfaces and support the design and interpretation of scattering experiments.

**TOC Graphic:** 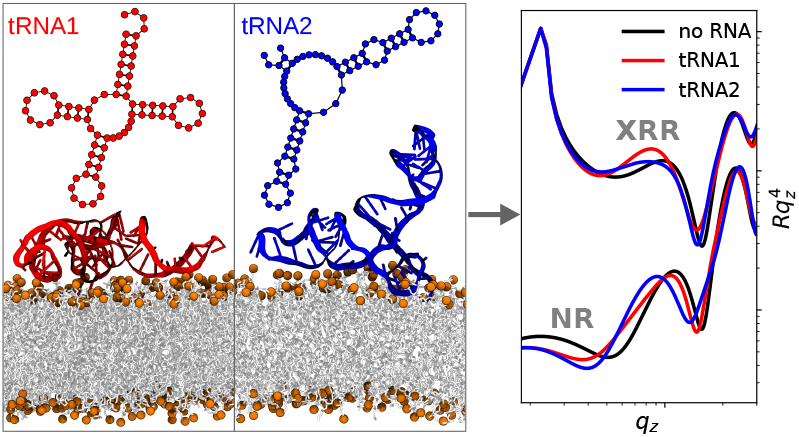

## Introduction

Understanding how RNA interacts with lipid assemblies at lipid/water interfaces is essential for elucidating biological processes such as viral entry, membrane fusion, and RNA delivery via lipid nanoparticles.^1–4^ The lipid environment can influence RNA conformation, folding, and degradation, all of which are key factors in determining the efficacy of RNA-based therapeutics .^2,3,5–10^

Despite their biomedical significance, RNA-lipid interactions remain challenging to characterize at the molecular level under physiologically relevant conditions. XRR and NR are powerful tools for probing such interactions, providing sub-nanometer structural information normal to the interface. Both techniques have been widely applied to study supported lipid bilayers and monolayers at the air/water interface,^11–15^ as well as interactions with ions and biomolecules including peptides, and nucleic acids.^16–23^ Reflectivity measurements yield scattering length density (SLD) profiles across the interface encoding the spatial distributions of molecular components, such as lipid tails, head groups, and RNA, enabling inference of orientation, insertion, and adsorption geometry.^15,18,24^

However, the interpretation of reflectivity data is nontrivial. The scattering profile cannot be directly inverted into atomic positions, and traditional fitting with slab models lacks molecular details.^25^ As a result, the structure of RNA at lipid/water interfaces remains poorly understood. Molecular dynamics (MD) and coarse-grained (CG) simulations address this gap by providing molecular representations of RNA-lipid systems. ^26,27^ MD simulations reveal lipid packing, RNA binding modes, and hydration structure, and allow the calculation of SLD profiles for direct and model free comparison with experimental scattering data. ^19,20,28^ For example, simulations combined with NR were used to assess the distribution of ionizable lipids in bilayers and the quality of different simulation models.^20^ Integration with X-ray scattering and fluorescence enabled the determination of lipid packing and pH-dependent protonation states in lipid monolayers^19^ and the structure and composition of adsorbed mRNA layers.^23^ Likewise, combination of SAXS and MD simulations successfully reproduced lyotropic phases with correct mesophase symmetries, water content, and pH-induced structural transitions.^4,29,30^ Similarly, CG simulations showed good agreement with NR profiles of lipid bilayers^31^ and elucidated pH-dependent interaction between RNA and ionizable lipids,^32^ further highlighting the benefit of joint simulation and experimental scattering approaches. In addition to a molecular interpretation of scattering signals, simulations can also guide experimental design by enabling *a priori* prediction of reflectivity curves, optimizing parameters like deuteration and sample composition to improve signal-to-noise and reduce beam time.

In this study, we use Martini CG simulations to investigate the interactions of singlestranded (ssRNA), double-stranded RNA (dsRNA), and tRNA with neutral DOPC and cationic DOTAP lipid bilayers. Subsequently, we address the question how RNA adsorption and RNA secondary structure can be detected experimentally via surface scattering techniques including neutron and X-ray reflectivity. We compute the reflectivity profiles to assess how RNA structure, concentration, lipid composition, and solvent contrast influence the corresponding scattering signals.

## Materials and Methods

### System Setup and Simulation Details

We simulated DOPC, DOTAP and mixed DOPC/DOTAP bilayers in the presence of single-stranded RNA (ssRNA), doublestranded RNA (dsRNA), and transfer RNA (tRNA) molecules. For systems containing ssRNA or dsRNA, five DOPC:DOTAP molar ratios were used: 1:0, 3:1, 1:1, 1:3, and 0:1, resulting in five distinct bilayer simulation setups. For tRNA-containing systems, only 100% DOPC and 100% DOTAP bilayers were simulated.

The lipids were described using the Martini force field (version 2.1), and RNA molecules were modeled using the corresponding Martini RNA force field.^27,33^ The Martini 2.1 force field was employed to ensure consistency with trajectories generated at the start of the project, prior to the publication of Martini 3. The initial configurations were generated using the insane.py script. ^34^

To construct a CG RNA structure, a doublestranded atomistic model was first gener-ted using the fd helix.c code from Ref.,^35^ which was subsequently coarse-grained using the martinize-nucleotide.py script. ssRNA was obtained by removing the complementary strand from the double-stranded structure. The RNA molecules were positioned approximately 2 nm away from the bilayer surface. All systems were solvated with CG water, including 10% anti-freeze water beads. CG sodium and chloride ions were added to achieve a physiological salt concentration of 150 mM and to ensure overall charge neutrality.

The ssRNA and dsRNA systems contained 100 lipids per monolayer, while the tRNA systems included 324 lipids per monolayer. The ssRNA sequence was a polyU chain of seven nucleotides, whereas the dsRNA comprised the sequence UCUUCUACUU and its complementary strand, which was used in a previous experimental study on RNA-lipid interactions.^36^

The initial configurations were energy-minimized using the steepest descent algorithm. This was followed by a 100 ns equilibration phase in the NPT ensemble, employing the Berendsen barostat and thermostat to maintain a pressure of 1.0 bar and a temperature of 323 K, respectively. The elevated temperature was used to enhance conformational sampling of tRNA on the bilayer surface.

During the production run, the temperature was maintained at 323 K using the stochastic velocity rescaling thermostat with a time constant of 1.0 ps,^37^ and the pressure was controlled semi-isotropically at 1.0 bar using the Parrinello-Rahman barostat with a time constant of 12.0 ps. ^38^ In all cases, the RNA molecule was restrained to the initial 3D conformation using an elastic network with a force constant of 1000 kJnm^*−*2^mol^*−*1^.

All simulations were performed using the GROMACS software package (versions 2018 and 2021).^39^ Van-der-Waals interactions were truncated at 1.1 nm and smoothly shifted to zero. Long-range electrostatic interactions were treated using the reaction-field method with a 1.1 nm cutoff and a relative dielectric constant of 15 which are typical settings for Martini force fields.^26^ The equations of motion were integrated with a time step of 10 fs. Systems with ssRNA and dsRNA were simulated for 5 *µ*s, while systems containing tRNA in various conformations were simulated for 1–3 *µ*s each. All analyses were performed using built-in GROMACS tools or custom Python scripts based on the MDAnalysis package.^40^ Visualizations were generated using VMD software.^41^

### All-atom Simulations

We performed all-atom simulation of a pure DOPC bilayer with 100 lipids per leaflet using the Gromacs simulation package. The initial setup was created using the Charmm-gui web-server. ^42^ The bilayer was hydrated with 50 water molecules per lipid and 150 mM of NaCl was also added. The lipid molecules were described using the Amber lipid14, ^43^ water with TIP3P^44^ and the ions with the Mamatklov-Schwierz force field^45^ which was optimized for TIP3P water. The structure obtained from Charmm-gui was energy minimized with a gradient descent algorithm followed by 1.0 ns equilibration with semi-isotropic C-rescale barostat^46^ with a time constant of 5.0 ps and the stochastic velocity rescaling thermostat^37^ with a time constant of 1.0 ps. The temperature was maintained at 300 K and pressure at 1.0 bar. The production run was carried out for 200 ns with Parrinello-Rahman barostat^38^ with 5.0 ps time constant. The Van-der-Waals interactions were cut-off and shifted to zero at 1.2 nm. Electrostatic interactions were evaluated using the particle mesh Ewald (PME) method^47^ with a cut-off of 1.2 nm. The electron density profile was calculated after excluding the initial 50 ns of the trajectory for equilibration.

### X-ray and Neutron SLD Profiles from CG Simulations

The neutron and X-ray SLD profiles, denoted as *ρ*_*n*_(*z*) and *ρ*_x_(*z*), respectively, along the bilayer normal from simulations were obtained by dividing the simulation box into small slices as:

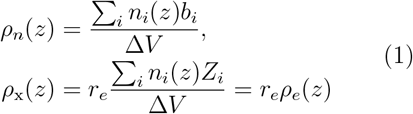

where *n*_*i*_(*z*) is the number of atoms of type *i* in a slice of volume Δ*V* centered at *z, b*_*i*_ is the coherent neutron scattering length, *Z*_*i*_ is the number of electrons of atom *i, r*_*e*_ = 2.82 × 10^*−*6^ Å is the classical electron radius and *ρ*_*e*_(*z*) is the transverse electron density profile. We employed two distinct methods to assign the scattering length, *b*_*i*_ or the number of electrons, *Z*_*i*_, to the CG Martini beads:

1. **Direct calculation from CG simulations** In this method, *b*_*i*_ or *Z*_*i*_ was directly assigned to the CG beads and Eqn. 1 was used to calculate *ρ*_*n*_(*z*) or *ρ*_x_(*z*). The *b*_*i*_, *Z*_*i*_ values for the beads were determined using the Martini mapping file provided with the force field for each molecule. Based on this, the *b*_*i*_, *Z*_*i*_ value of a bead was computed as the sum of the *b*_*i*_ or *Z*_*i*_ values of all atoms mapped into it, weighted by their fractional contributions. An example mapping for DOPC is shown in Table S1, where the contribution of each atom to a CG bead is specified. This method is highly efficient as it eliminates the need for backmapping and reduces disk space usage. Our scripts that automate this task are freely available at https://git.rz.uni-augsburg.de/cbio-gitpub.
2. **Backmapping and MD Equilibration:** In this method, approximately 2000 coarse-grained frames were backmapped to all-atom resolution using the backward.py script. ^48^

Each frame was projected to all-atom resolution, followed by energy minimization and a 2.0 ps MD simulation. These backmapped and equilibrated frames were then analyzed using a custom script to compute *ρ*_*n*_(*z*) or *ρ*_x_(*z*) according to Eqn. 1, using known atomic *b*_*i*_ or *Z*_*i*_ values. This method is computationally more expensive and requires substantial disk storage.

### Undulation Corrections

NR and XRR curves, as well as scattering form factors, are determined by the neutron or Xray SLDs along the bilayer normal (Eqn.1). For small bilayer systems (∼100 lipids/monolayer), these density profiles can be obtained by binning the system along the transverse direction, assuming the bilayer to be flat. However, in simulations of larger lipid patches, membrane undulations become significant, and simple binning under the flat-surface assumption produces a smeared-out density profile (Figure S8). Such undulations also exist in experiments at various length scales, but in supported bilayers or in monolayers at the air-water interface they may be suppressed either due to attachment to the planar substrate or the high surface tension of the interface, respectively. Therefore, to enable direct comparison between experiments and simulations, the respective undulations must be removed.^49^ To correct for the undulations in the simulations, we employed the method developed by Braun et al. ^49^ For each frame in the trajectory, this method constructs an undulating reference surface using the Fourier coefficients of selected atom groups in the bilayer, followed by binning with respect to this dynamically defined surface. Further details and evidence of improved agreement with experimental results are provided in the Supporting Information, Section 4, and Figure S8. Our implementation is freely available at https://git.rz.uniaugsburg.de/cbio-gitpub. Note that in simulations of very large lipid bilayer systems, undulations can also arise from neighbor list update artifacts.^50^ However, the systems studied here are well below the size at which such effects become significant.

### NR and XRR Curves

The NR and XRR curves were obtained from *ρ*_*n*_(*z*) and *ρ*_x_(*z*) using the Abeles matrix formalism,^51^ as implemented in the Refnx Python package.^52^ The SLD profiles from simulations were discretized into layers of constant values, each 1 Å thick. NR and XRR curves were obtained for two experimental setups (i) silicon– supported lipid bilayers and (ii) lipid monolayers at the air/water interface. In both cases, we used MD simulations of free lipid bilayers in water (see section *System Setup and Simulation Details*). For supported bilayers on silicon substrates, the silicon not present in the simulations was incorporated following our previous work.^20^ Specifically, the SLD values for silicon and silicon dioxide layers were set to known values, with the SiO_2_ layer thickness fixed at 10 Å, which is typical for such experiments. ^20^ The Si– SiO_2_ density profile was merged with the total lipid density profile at the point below (for NR) or above (for XRR) where the lipid density becomes zero, yielding the complete density profile. NR and XRR reflectivity profiles were then calculated from this profile using Refnx. Although the solid supports of bilayers can, in principle, influence lipid structure, the contribution to the reflectivity signal is expected to be small. ^20^ The major contributions arising from the solid support are changes in the interfacial water fraction and the presence of water patches, which can be explicitly modeled when experimental reflectivity data are available to constrain the model. ^20,31^

In the second setup, the monolayer was constructed by selecting a single leaflet from the bilayer. Given the area per lipid in our simulations (∼69.4 Å^2^), this monolayer setups corresponds to experimental conditions with a surface pressure of approximately 15–20 mN/m.^53^ Under these conditions, the resulting lipid packing and hydration are expected to be representative of the air/water interface experiments, enabling a meaningful comparison between simulations and experimental observations of RNA adsorption.

### tRNA Secondary and Tertiary Structure Prediction

We used a tRNA molecule with the sequence GCGGAUUUAGCUCAGUUGGGAGAGCGCCAGACUGAAGA UCUGGAGGUCCUGUGUUCGAUCCACAGAAUUCGCACCA.

The secondary structure of the sequence was predicted using the RNAfold package.^54^ The tRNA exhibited a minimum free energy of -22.40 kcal/mol. The minimum energy structure, along with the nine closest structures within 18 kcal/mol and separated by at least 1.5 kcal/mol, were used in the simulations (Figure S9). The RNAComposer webserver was used to generate an atomistic 3D structure for each secondary structure,^55^ which was then coarse-grained using the martinize nucleotide.py script.

## Results and Discussion

In the following sections, we discuss the interactions of ssRNA and dsRNA with lipid bilayers focusing on the influence of RNA structure and lipid charge. Subsequently, we evaluate the accuracy of the coarse-grained model in reproducing experimental X-ray scattering form factors using different backmapping strategies. Finally, we compute how RNA adsorption affects neutron and X-ray reflectivity curves of supported bilayers and monolayers at the air/water interface. We systematically analyze the effects of contrast conditions, lipid-to-nucleotide ratio, lipid composition, and RNA secondary structure.

### Interactions of ssRNA and dsRNA with Lipid Layers of Increasing Charge

We simulated ssRNA and dsRNA on bilayers with increasing fractions of cationic DOTAP lipids using CG simulations. In all cases, the RNA adsorbs at the lipid/water interface and displays distinct interaction modes depending on base-pairing and membrane compositions. For ssRNA, the nucleobases are embedded near the hydrophobic lipid tails and the sugar phosphate backbone is associated with polar head groups (Figure 1A–C), consistent with previous studies. ^5,56^ As the cationic lipid content increases, electrostatic attraction strengthens ss-RNA adsorption, resulting in a more narrow distribution close to the bilayer surface. dsRNA interacts more weakly with neutral bilayers due to base-pairing that shields hydrophobic regions (Figure 1A). On the charged bilayers, dsRNA adsorbs due to the electrostatic attraction but adopts a more diverse range of orientations, particularly at intermediate charge fractions leading to broader distributions (Figure 1E–G and S1).

**Figure 1.**
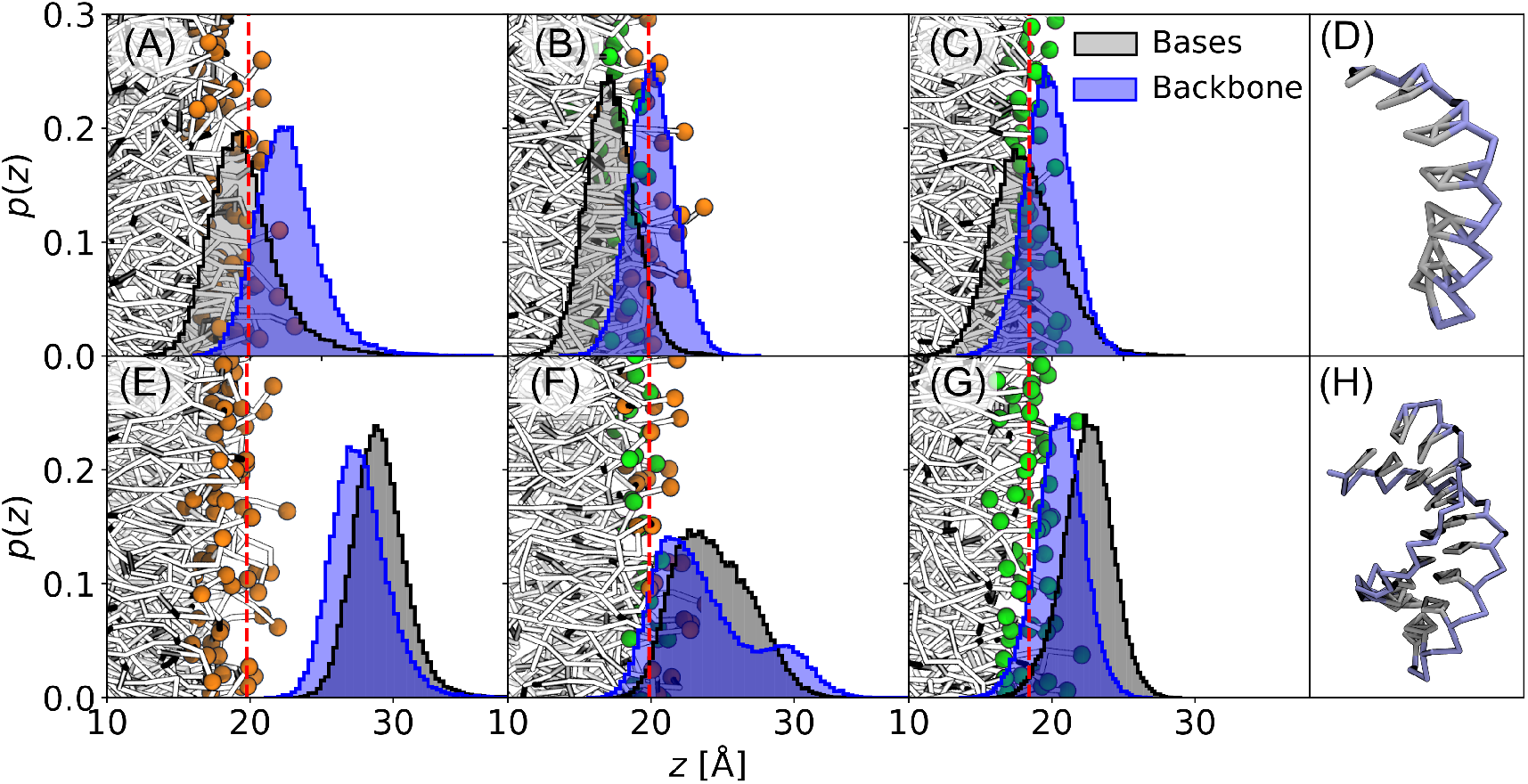
Distribution of ssRNA and dsRNA at the lipid/water interface. Normalized probability distrivution *p*(*z*) of finding a nucleobase (gray) or backbone phosphate (blue) along the membrane normal for ssRNA (top) and dsRNA (bottom). (A) *p*(*z*) for ssRNA on a pure DOPC bilayer. Orange spheres indicate the DOPC choline headgroups. (B) *p*(*z*) for ssRNA on a mixed DOPC/DOTAP bilayer (1:1 molar ratio). Green spheres represent the DOTAP choline headgroups. (C) *p*(*z*) for ssRNA on a pure DOTAP bilayer. (D) CG model of ssRNA, with nucleobase and backbone regions colored gray and blue, respectively. (E–G) *p*(*z*) for dsRNA on the bilayer with same composition as above. (H) CG structure of dsRNA, with color scheme identical to (D). Red dashed vertical lines denote the average position of the lipid choline headgroups.

These results align with experimental and all-atom simulation data showing weaker dsRNA–membrane interactions compared to ssRNA,^12,57^ supporting that the CG Martini model captures the key electrostatic and hydrophobic contributions to RNA–bilayer binding. Recent all-atom MD simulations^58^ also reported weaker dsRNA interactions with lipid bilayers compared to ssRNA, in agreement with our findings. In those studies, the weaker interaction was attributed to hydrogen bond formation between RNA base pairs and lipid head groups which are not explicitly represented in the current Martini CG model but are implicitly embedded in the Lennard-Jones and electrostatic interaction parameters. Accordingly, the interactions observed in our simulations are interpreted in terms of effective electrostatic and hydrophobic driving forces.

Finally, the interaction between RNA and lipids also affects the dynamics of both components. The diffusion coefficient of the lipids in the leaflet in contact with RNA is consistently lower than that of the opposite leaflet as expected. The difference between the leaflets increases as the fraction of cationic DOTAP lipids increases (Figure S2G, H). The diffusion coefficient of ssRNA and dsRNA on mixed neutral and cationic bilayers is slightly lower compared to fully charged or fully neutral bilayers (Figure S2G, H). This trend in diffusion coefficient correlates inversely with the increased surface roughness of mixed-composition bilayers relative to pure neutral or cationic bilayers. (Figure S3-S4).

### Comparison of Different Backmap-ping Procedures to Reproduce Experimental Form Factors from Xray Scattering

Next, we analyzed how well the CG model reproduces the experimental scattering data of lipid bilayers. To this end, we compare the X-ray scattering form factors calculated from DOPC bilayer simulations with experimental form factors obtained from multilamellar lipid stacks.^28^The X-ray scattering form factor (|*F*(*q*_*z*_)|) is a standard quantity used to validate the structure of lipid bilayers^28,43,59^ and probes the transverse electron density profile. The form factor can be obtained directly from simulations via a Fourier transform of the electron density profiles.

The electron density profiles and corresponding form factors were computed using several CG-to-all-atom reconstruction protocols. We first applied the direct method, where form factors are computed directly from the CG electron density. This approach, while computationally efficient, leads to inaccurate bilayer core densities, particularly for long-tail lipids such as DOPC (Figure 2A). This artifact arises because the Martini force field maps lipid tails of varying length to the same number of beads (e.g., four beads for both DOPC and DPPC), concentrating more electrons in the terminal beads of longer lipids and artificially increasing the central density. However, for lipids with relatively shorter tails like DPPC, such artifacts do not arise (Figure S6).

**Figure 2.**
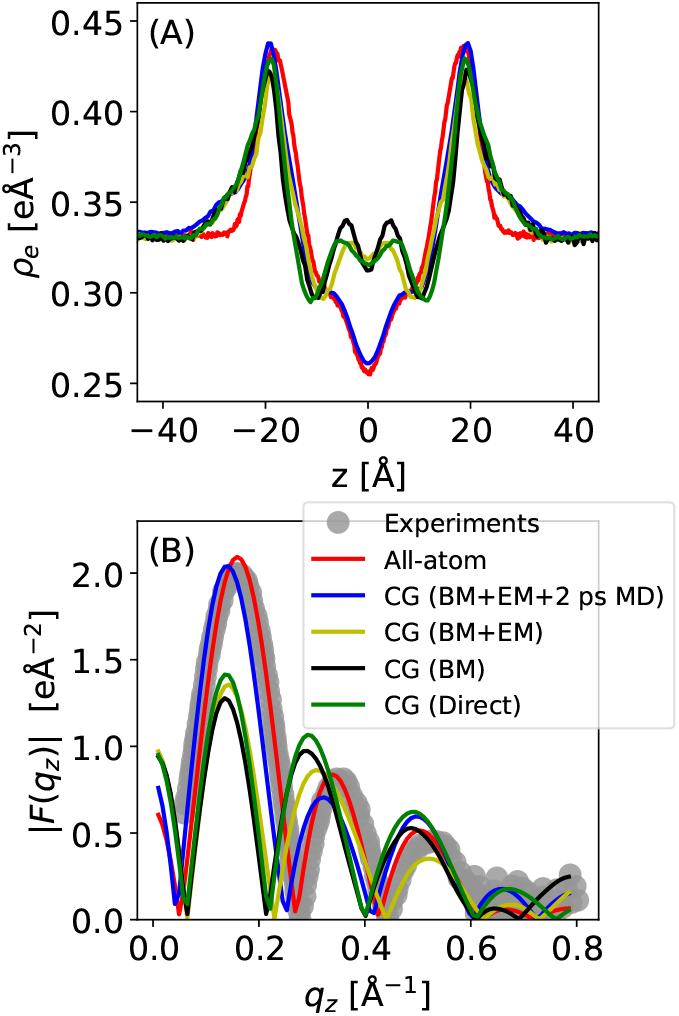
Comparison of different backmapping strategies to reproduce experimental X-ray scattering data for a DOPC bilayer. (A) Electron density *ρ*_*e*_(*z*) obtained from all-atom MD and CG simulations with different backmapping protocols. Four CG-to-all-atom backmapping protocolls were assessed: (i) direct CG (no backmapping), backmapped to all-atom resolution (BM), backmapped followed by energy minimization (BM+EM), and (iv) backmapped with energy minimization followed by a short 2 ps MD equilibration (BM+EM+2ps MD). (B) Corresponding X-ray scattering form factors derived from each approach, compared to experimental data from Ref. ^28^

To improve the agreement with experiments, we employed the standard backmapping pipeline^48^ and evaluated three progressively refined backmapping strategies. In the BM (backmapped) protocol, coarse-grained frames were projected to all-atom resolution without relaxation. The resulting electron density profiles still showed central density artifacts comparable to the direct CG method. The BM+EM (backmapped with energy minimization) approach applied two sequential minimization, first considering bonded terms only, followed by inclusion of non-bonded interactions. This procedure improved stability but did not significantly alter the density profile. In the final BM+EM+2 ps MD protocol, which adds a short 2 ps MD equilibration with gradually increasing time steps (0.1 to 2.0 fs), the electron density profile converges to the shape and peak-to-peak spacing observed in all-atom simulations and experiments ^28^ (Figure 2B). The observed minor deviations at the water/lipid interface, where CG simulations show slightly elevated water density, are consistent with the known reduced water penetration in Martini models due to the larger size of CG water beads.^60^ Among all tested methods, only the BM+EM+2 ps MD approach reproduces both the bilayer thickness and the overall shape of the electron density distribution. We therefore adopt the BM+EM+2 ps MD procedure for all subsequent calculations of X-ray and neutron scattering length density profiles and reflectivity signals.

### NR and XRR for tRNA at Lipid/Water Interfaces

In the following, we discuss how RNA adsorption influences NR and XRR signals from supported bilayers and monolayers at the air/water interface. We evaluate the effects of contrast conditions, lipid-to-nucleotide ratio (*L/N*), lipid charge, and RNA conformation. The question of RNA conformation is motivated by experimental studies showing that RNA adsorption is highly sensitive to features such as base-pairing and single-stranded regions, ^5,56^ as well as by previous applications of reflectivity to resolve protein and peptide conformations at bilayer surfaces.^21,22^ To investigate the impact of RNA conformation we used nine distinct tRNA secondary structures (Figure S9). In the simulations, these structures adopt conformation-specific orientations on the neutral DOPC and cationic DOTAP bilayers and show distinct binding geometries and distributions (Figure 3 and Figure S12). In some cases (e.g. tRNA9 on DOPC), we observe detachment and re-adsorption of tRNA, indicating that the tRNA explores different binding modes on the *µ*s timescale. The adsorption behavior of tRNA is consistent with the behavior of ssRNA and dsRNA discussed above. In most cases, tRNA adsorbs to neutral bilayer via the single-stranded regions whereas on cationic bilayers, it interacts via both single and double stranded regions (Figure 3, Figure S10S11). For each of the systems shown in Figure 3, three additional 2 *µs* replicas with different initial tRNA orientations relative to the bilayer were simulated (Figures S14-S17). Most replicas reproduced the binding behavior shown in Figure 3; however, some exhibited alternative binding modes, highlighting the challenge of exhaustively sampling all possible configurations even in coarse-grained simulations. However, the secondary structure binding patterns remain consistent in all cases.

**Figure 3.**
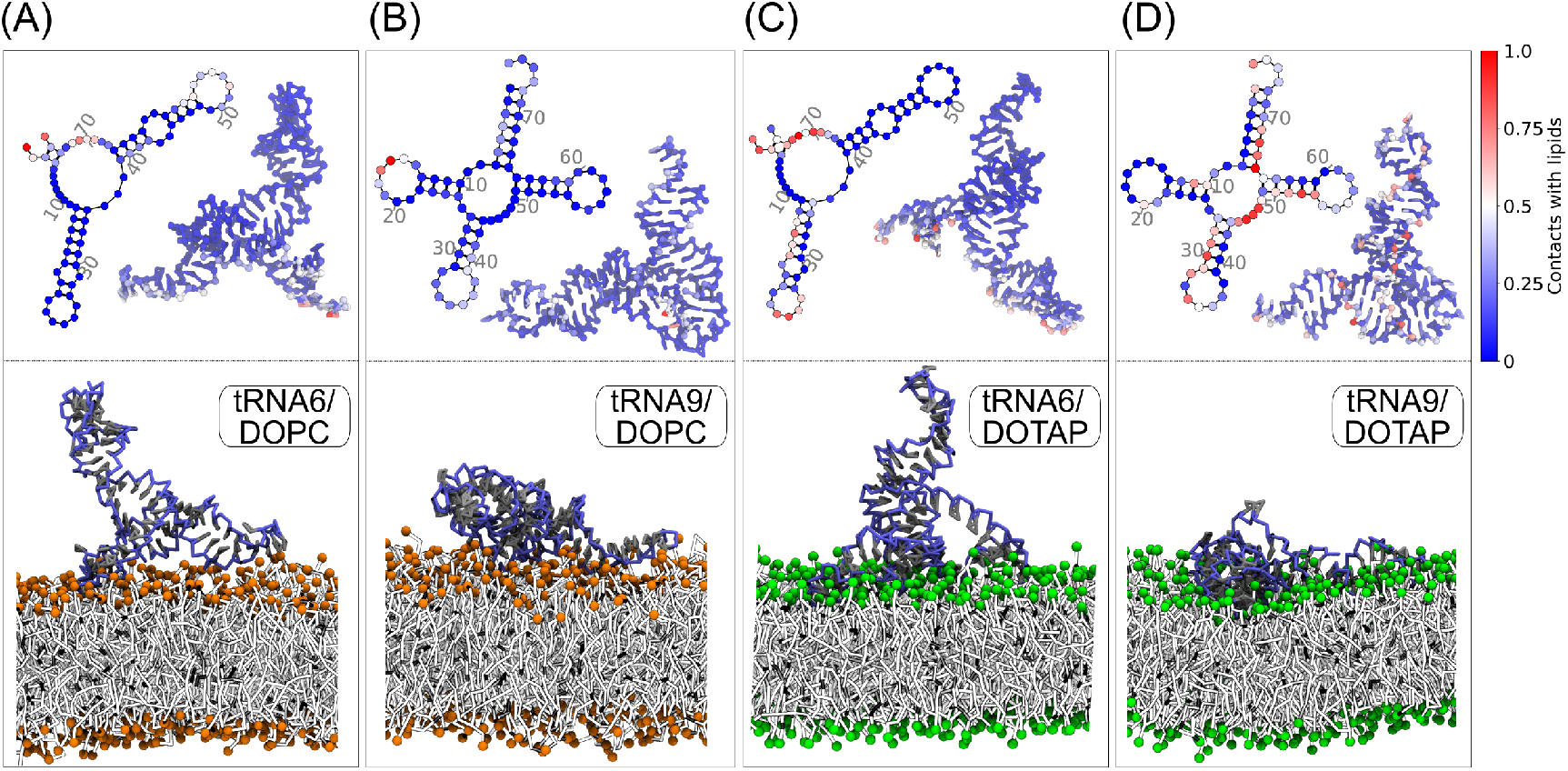
Simulation snapshots of two tRNA structures interacting with neutral and charged lipid bilayers after 3 *µ*s of simulation time. (A) tRNA6 on a neutral DOPC bilayer. (B) tRNA9 on a DOPC bilayer. (C) tRNA6 on a positively charged DOTAP bilayer. (D) tRNA9 on a DOTAP bilayer. tRNA6 and tRNA9 are the conformations which exhibited the largest differences in NR and XRR profiles. Top: tRNA secondary and tertiary structures with colors indicating membrane contacts. Red indicates high contact levels, blue low contact levels. For each structure, contact values are normalized to their respective maximum. All other tRNA structures are shown in Figure S10-S11). Bottom: Simulation snapshots of the two tRNA conformations. tRNA backbones are depicted in blue, nucleobases in gray. Choline headgroups of DOPC and DOTAP lipids are shown as orange and green spheres, respectively.

We considered two experimentally relevant setups: (i) supported bilayers on silicon substrates, ^20^ and (ii) monolayers at the air/water interface.^19,23^ NR profiles were calculated for four different deuteration conditions: (i) non-deuterated solvent (H_2_O) with regular non-deuterated lipid (hLipid), (ii) deuterated solvent (D_2_O) with hLipid, (iii) H_2_O with deuterated lipid (dLipid), and (iv) D_2_O with dLipid. For the monolayer case, we used null contrast conditions (∼ 92% H_2_O and 8.0% D_2_O) instead of pure H_2_O, as is typically done in such exper-iments.^61,62^

We begin with the NR results for supported bilayers and monolayers (Figure 4A,D). For *L/N* = 1.5, the clearest differences in the reflectivity signal were observed for supported bilayers using the hLipid/H_2_O contrast and for monolayers under null contrast (Figure S11 and S18). Under these conditions, differences between the systems with and without RNA, and between the two selected tRNA structures are visible (Figure 4C,F). For bilayers, these differences are most pronounced in the *q*_*z*_ range 0.025–0.16 Å^*−*1^ (Figure 4C). For monolayers they appear for *q*_*z*_ *>* 0.025 Å^*−*1^ (Figure 4F). Notably, RNA adsorption shifts the first reflectivity minimum toward lower *q*_*z*_, consistent with an increase in total thickness due to the RNA adsorption.^23^

**Figure 4.**
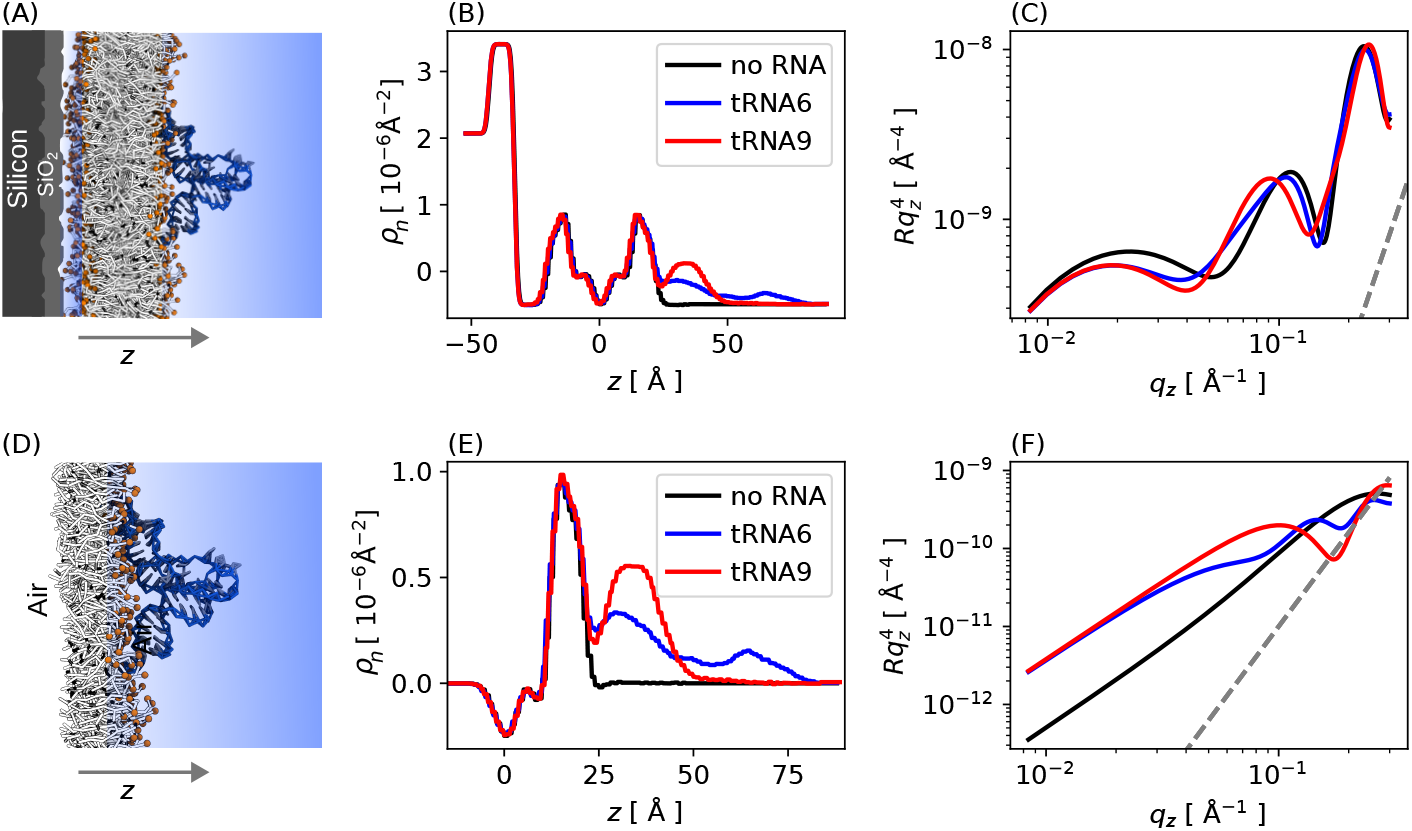
NR curves of RNA interacting with supported bilayers (top) and monolayers (bottom). (A) Solid-supported bilayer setup: Silicon/SiO_2_ substrate, DOPC bilayer, RNA, and water. The neutron beam is incident from the left (*z <* 0). (B) Neutron SLD *ρ*_*n*_ for tRNA6 and tRNA9 on DOPC bilayers corresponding to setup (A). (C) NR curves for the systems shown in (B). Monolayer setup: Air, DOPC monolayer, RNA, and water. (E) Corresponding neutron SLD profiles for tRNA6 and tRNA9 interacting with DOPC monolayers. (F) Monolayer systems in (E). The dashed gray lines in (C) and (F) represent the background noise level, defined as 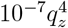For an explicit noise model for reflectivity see SI Section 2. In all cases, the lipid-to-nucleotide ratio is *L/N* = 1.5. For the bilayer setup hLipid/H_2_O contrast is used whereas for the monolayer null contrast (∼ 92% H_2_O and 8.0% D_2_O) is used in place of H_2_O.

As expected, under deuteration conditions involving D_2_O and/or dLipid, such differences are difficult to resolve (Figure S11 and S12). In these cases, the RNA contribution to the overall SLD is small compared to that of the deuterated components, resulting in reflectivity curves that are largely insensitive to RNA presence or conformation.

We now turn to X-ray reflectivity (XRR) as a complementary approach to probe RNA ad-sorption (Figure 5A, D). For *L/N* = 1.5, differences in reflectivity curves are observed between systems with and without RNA, and between two selected tRNA structures (Figure 5C, F) but are slightly less pronounced compared to NR. The differences appear only in the *q*_*z*_ range of 0.03–0.12 Å^*−*1^ for bilayers and up to 0.2 Å^*−*1^ for monolayers.

**Figure 5.**
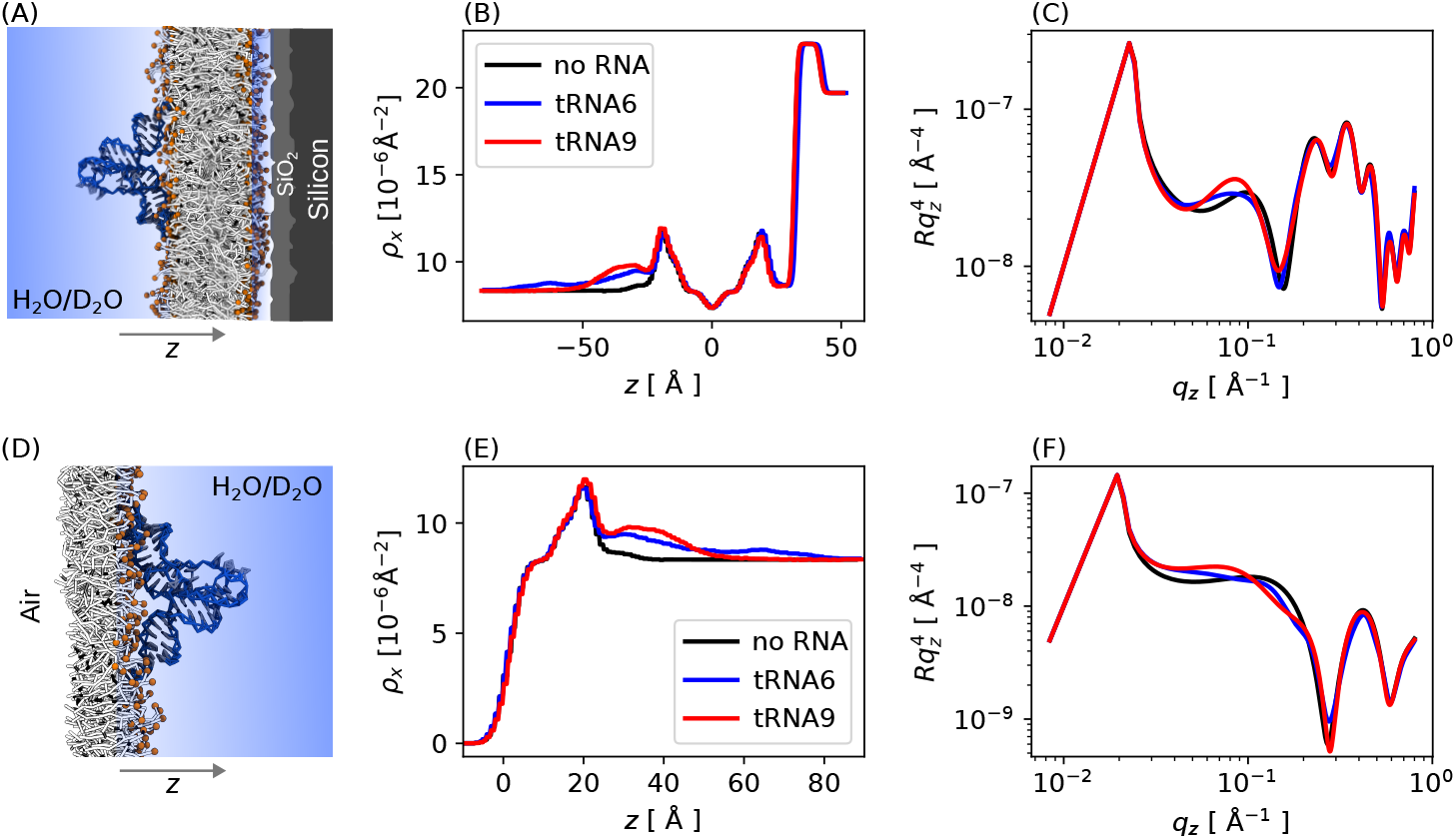
XRR curves of RNA interacting with solid supported bilayers (top) and monolayers (bottom). (A) Solid-supported bilayer setup. The X-ray beam is incident from the left (*z <* 0). (B) X-ray SLD *ρ*_x_ for tRNA6 and tRNA9 on DOPC bilayers corresponding to setup (A). (C) X-ray reflectivity curves for the systems shown in (B). (D) Schematic of the monolayer setup: air, DOPC monolayer, RNA, and water. (E) Corresponding *ρ*_x_ for tRNA6 and tRNA9 interacting with DOPC monolayers. (F) X-ray reflectivity data for monolayer systems in (D). The line depicting the background noise level is omitted here since it is much lower than the signal. In all cases, the lipid-to-nucleotide ratio is *L/N* = 1.5.

Finally, we provide an overview of the factors that influence NR and XRR profiles: lipid-to-nucleotide ratio, lipid charge, and RNA conformation Figure 6. We focus on the NR signals at the supported bilayers (top) and XRR signals (bottom) for the monolayers at the air/water interface (see supporting information for all other setups).

**Figure 6.**
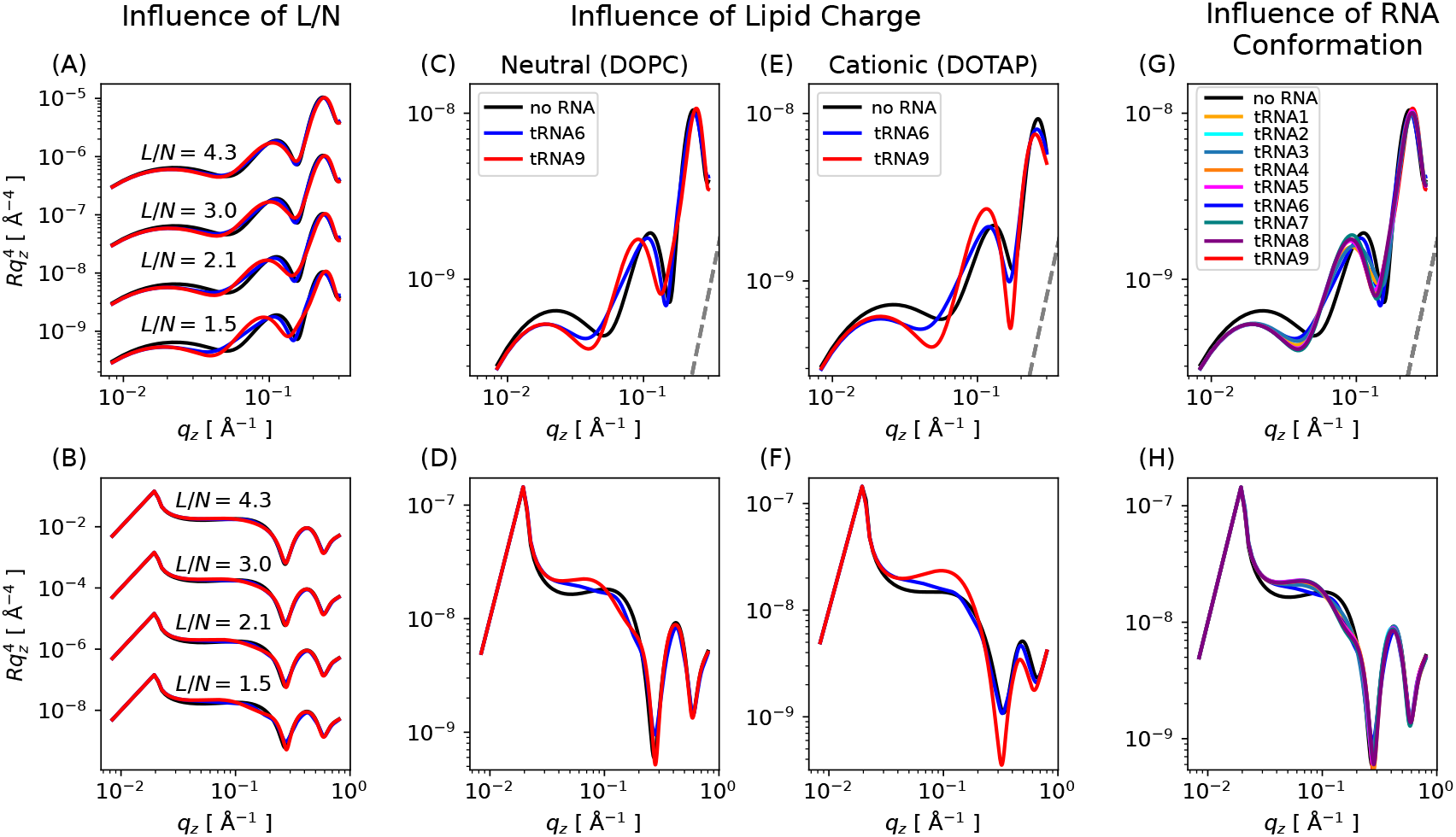
Influence of L/N ratio, lipid charge and RNA conformation on NR curves for solid-supported bilayers (top) and XRR curves for monolayers at the air/water interface (bottom). Left: Influence of *L/N* ratio. (A) NR from a solid-supported DOPC bilayer without RNA, with tRNA6 and tRNA9. (B) XRR from a DOPC monolayer at the air/water interface with same tRNA variants. Middle: Influence of lipid charge. (C, E) NR from neutral DOPC and positively charged DOTAP bilayers without RNA, with tRNA6 and tRNA9. (D, F) XRR from neutral DOPC and positively charged monolayers with the same tRNA variants. Right: Influence of tRNA secondary structure. (G) NR from DOPC bilayers interacting with 9 different tRNA structures (Figure S9). (H) XRR from DOPC monolayers for same tRNA variants. Dashed gray lines indicate the background noise threshold, defined as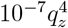. For XRR this noise is omitted since it is much lower that the signal. It is also omitted in (A) for clarity, where all signals are well above the noise. In panels (C–H), the ratio is fixed at *L/N* = 1.5. hLipid/H_2_O contrast was used for the neutron reflectivity curves.

The lipid-to-nucleotide *L/N* ratio is a key determinant of detectability. We investigated this by changing the *L/N* ratio from 4.3 to 1.5 (see supporting information section 9). Note that these ratios are similar to the ratio used in lipid nanoparticle (LNP) formulations.^63^

For NR, differences between pure lipid systems and with two different RNA conformations are clearly resolved only below *L/N <* 3.0 (Figure 6A). XRR can still detect the presence of RNA at this threshold but shows limited sensitivity to different RNA conformation (Figure 6B).

Lipid charge plays a pivotal role in RNA-lipid interactions. In systems containing cationic DOTAP lipids, both NR and XRR profiles exhibit more pronounced differences, which can be attributed to enhanced electrostatic interactions between the cationic lipids and anionic RNA molecules (Figure 6C–F).

The impact of RNA structure is further illustrated by comparing nine different tRNA variants (Figure 6G–H). While certain conformations (e.g., tRNA6 and tRNA9) lead to distinctly different reflectivity curves, others produce nearly indistinguishable NR and XRR signals.

Collectively, these findings underscore the value of NR and XRR as mutually complementary techniques for probing RNA-lipid interactions at lipid/water interfaces. At sufficiently low lipid-to-nucleotide (*L/N*) ratios, both methods are sensitive enough to detect RNA adsorption at the interface. Still, forward models such as CG simulations are required for the molecular interpretation of the reflectivity curves.

## Conclusions

This study addresses the modeling of RNA at lipid/water interfaces using CG simulations, validation using experimental scattering data, and prediction of NR and XRR profiles directly from the simulations. We systematically explored how ssRNA and dsRNA, as well as structured tRNA, interact with lipid DOPC/DOTAP bilayers of increasing positive charge. Our results show that RNA secondary structure is a key determinant of RNA-lipid interactions at lipid/water interfaces, with single-stranded regions exhibiting stronger interfacial affinity than double-stranded segments. ssRNA-bilayer interactions are governed by hydrophobic interactions with exposed bases and electrostatic attraction between RNA backbone and cationic lipid headgroups. dsRNAbilayer interactions, constrained by base pairing, are weaker, particularly with neutral bilayers. These trends also hold for tRNA. Simulations of distinct tRNA conformations, representing different secondary structures, reveal that adsorption to neutral DOPC bilayers occurs primarily through exposed single-stranded regions, whereas in cationic DOTAP systems, adsorption is dominated by electrostatic interactions between the negatively charged tRNA backbone and the positively charged lipid headgroups.

To validate the CG bilayer model, we calculated and compared form factors from Xray scattering. The results show that, while CG simulations reproduce experimental data qualitatively, agreement improves substantially when CG structures are backmapped to atomistic resolution and refined by energy minimization and short MD simulations.

Using this validated approach, we simulated NR and XRR curves for RNA-decorated solid-supported bilayers and monolayers at the air/water interface. We show that both NR and XRR can detect RNA adsorption at lipid interfaces, especially under optimal contrast conditions and at low enough lipid-to-nucleotide ratios (*L/N <* 1.5). While reflectivity curves are sensitive to major RNA conformational changes, subtle structural differences will be challenging to resolve.

Systematic simulations can help to identify the experimental conditions best suited to resolve such subtle differences. Moreover, forward models such as the CG simulations presented here are expected to play an increasingly important role in the molecular interpretation of reflectivity data. Overall, our results demonstrate that combined simulation-scattering approaches are well suited to provide molecular insight into RNA adsorption and structure at lipid/water interfaces and to guide the de-sign of future scattering experiments on RNAmembrane systems.

## Acknowledgment

This work was supported by the German Federal Ministry of Education and Research through BMBF Project 05K18EZA within the framework of the Swedish-German research collaboration Röntgen-Ångström Cluster. The work was also supported by the Emmy Noether program of the Deutsche Forschungsgemeinschaft (DFG, German Research Foundation), number 315221747 and the research support program (Forschungspotenziale besser nutzen!) of the University of Augsburg. The authors gratefully acknowledge the scientific support and HPC resources provided by the Erlangen National High Performance Computing Center (NHR@FAU) of the Friedrich-AlexanderUniversität Erlangen-Nürnberg (FAU) under the NHR project b119ee and b253ee. We are grateful for the excellent support provided by the Max Planck Society. In particular, we are thankful to Gerhard Hummer at the Max Planck Institute for Biophysics, Frankfurt, Germany for his generous support as scientific host and for countless inspiring scientific discussions.

## Supporting Information Available

Noise model for reflectivity and data for XRR and NR calculations for all the nine tRNA secondary structures for different deuteration conditions and RNA surface concentration for both bilayer and monolayer setups, undulation correction procedure and Martini bead electron and neutron scattering factor assignment procedures. The codes to obtain undulation corrected densities from CG-Martini and allatom simulation trajectories are available at https://git.rz.uni-augsburg.de/cbio-gitpub.

