## supplementary files for "RNA at Lipid/Water Interfaces: Molecular Insights from Coarse-Grained Simulations and Reflectivity Data Predictions"

### Contents

**1 ssRNA and dsRNA on Lipid Bilayers** **S3**

1.1 RNA Orientation . . . . . . . . . . . . . . . . . . . . . . . . . . . . . S3

1.2 Mean Square Displacement and Diffusion Coefficient of RNA and Lipids . . S4

**2 Noise Model for Reflectivity** **S5**

**3 Assigning Neutron Scattering Factors /Electron /Mass to CG-Martini Beads** **S7**

**4 Undulation Correction** **S9**

**5 RNA Secondary Structure and Contact Frequency Maps** **S12**

**6 Additional Simulations for tRNA6 and tRNA9 on DOPC and DOTAP** **S16**

**7 NR Curves for tRNA at Bilayers and Monolayers** **S21**

7.1 tRNA/DOPC Systems . . . . . . . . . . . . . . . . . . . . . . . . . . . . S21

7.2 tRNA/DOTAP Systems . . . . . . . . . . . . . . . . . . . . . . . . . . . S23

**8 XRR for RNA at Bilayers and Monolayers** **S25**

8.1 tRNA/DOPC Systems . . . . . . . . . . . . . . . . . . . . . . . . . . . . S25

8.2 tRNA/DOTAP Systems . . . . . . . . . . . . . . . . . . . . . . . . . . . S26

**9 Lipid-to-Nucleotide Ratio Dependence of NR and XRR Signals** **S27**

**References** **S30**

### 1 ssRNA and dsRNA on Lipid Bilayers

#### 1.1 RNA Orientation

The orientation of RNA on the lipid bilayer is defined by the angle between the vector ($\vec{R}$) connecting the center of mass of two end nucleotides of the RNA and the bilayer normal ($\hat{n}$) and is given by,

$$\theta = \cos^{-1}\left(\frac{\vec{R}\cdot\hat{n}}{|\vec{R}|}\right) \tag{1}$$

$\theta = 90°$ corresponds to RNA in a flat orientation on the bilayer surface.

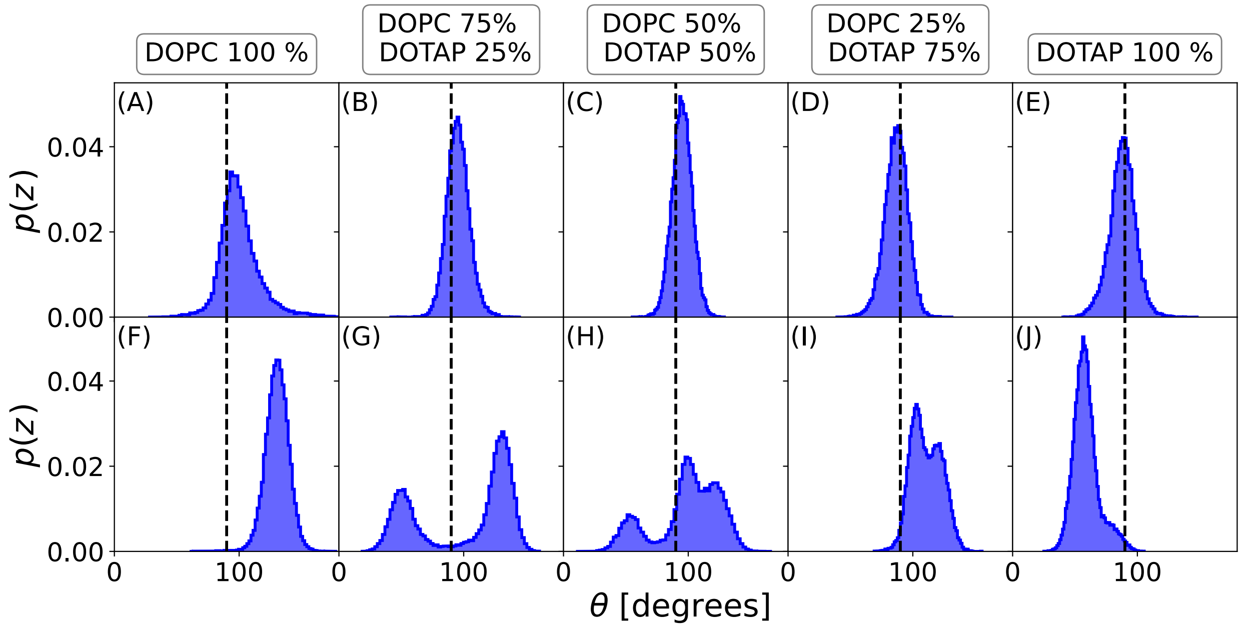


Figure S1: **RNA orientation with respect to the bilayer surface.** (A-E) ssRNA on bilayers with different ratios of DOPC and DOTAP. Each column corresponds to a particular membrane composition annotated on the top of each column. (F-J) dsRNA on bilayers. The vertical dashed line in each panel corresponds to $\theta = 90°$ i.e it lies in a flat configuration with its longer axis parallel to the bilayer surface.

#### 1.2 Mean Square Displacement and Diffusion Coefficient of RNA and Lipids

The dynamics of lipids and RNA can be characterized by the diffusion coefficient $D$. The diffusion coefficient in two dimensions (in the bilayer plane) is estimated from the mean square displacement (MSD) as

$$MSD(t) = 4Dt \tag{2}$$

Therefore, $D$ can be estimated by fitting the MSD profile in the diffusive regime (Figure S2 (A–C) and Figure S2 (E–G)).

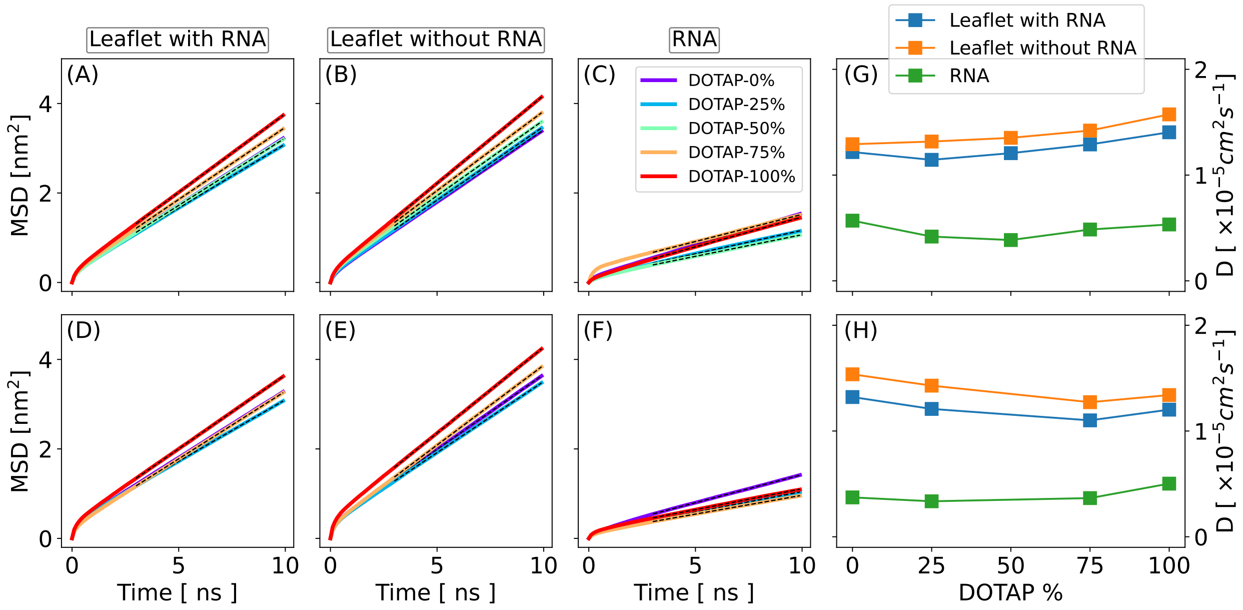


Figure S2: **Diffusion coefficients and mean squared displacements (MSDs).** Top panels: ssRNA–bilayer systems. Bottom panels: dsRNA–bilayer systems. (A) In-plane (2D) MSDs of the lipid center of mass in the bilayer leaflet where ssRNA is adsorbed for systems with different DOPC/DOTAP ratios. (B) MSDs of lipids in the opposite leaflet. The black dashed lines correspond to linear fits to Eqn. 2. (C) MSD of ssRNA. (D) Corresponding in-plane lipid and RNA diffusion coefficients calculated from Eq. 2. (E) MSDs of lipids in the leaflet containing dsRNA. (F) MSDs of lipids in the opposite leaflet. (G) MSD of dsRNA. (H) Corresponding diffusion coefficients of lipids and dsRNA.

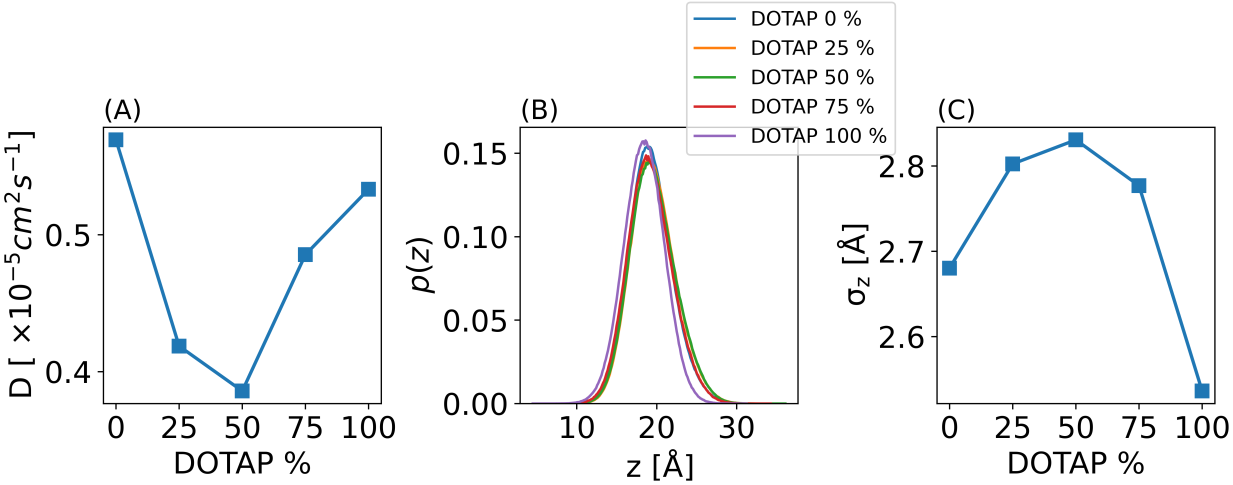


Figure S3: **Diffusion coefficients of ssRNA.** (A) Diffusion coefficients of ssRNA on bilayer with varying mixture of DOPC and DOTAP (same as green line in Figure S2G). (B) Distribution of lipid surface beads (NC3 and PO4) in z-direction from the bilayer center. (C) Surface roughness ($\sigma_z$) defined as the standard deviation of the distribution in B.

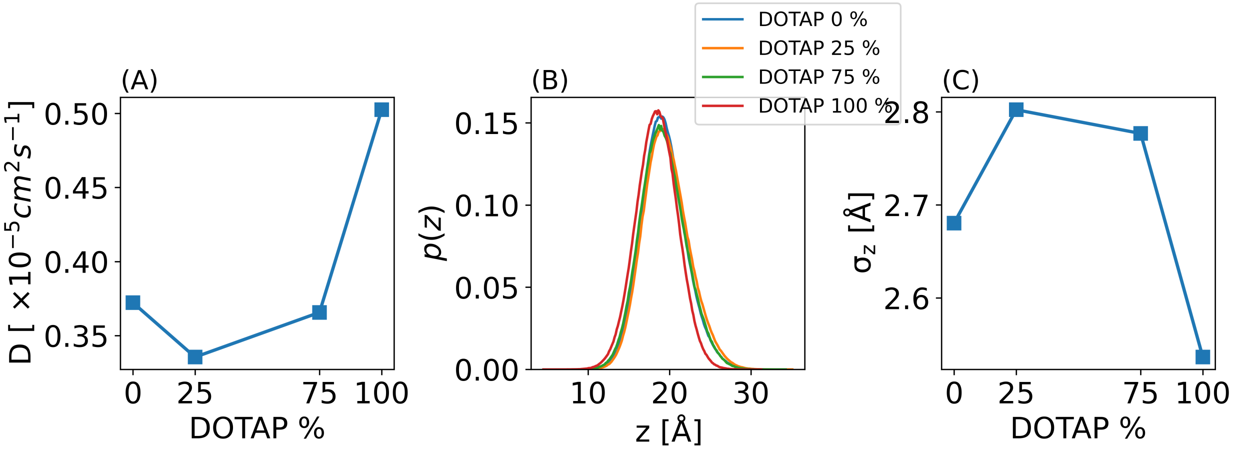


Figure S4: **Diffusion coefficients of dsRNA.** (A) Diffusion coefficients of dsRNA on bilayer with varying mixture of DOPC and DOTAP (same as green line in Figure S2H). (B) Distribution of lipid surface beads (NC3 and PO4) in z-direction from the bilayer center. (C) Surface roughness ($\sigma_z$) defined as the standard deviation of the distribution in B.

#### 2 Noise Model for Reflectivity

The total intensity measured in the experiments, $I_{meas}(q)$, contains contribution due to background scattering. Subtracting the background contribution, the reflected intensity from the sample is given by,

$$I_{refl}(q) = I_{meas.}(q) - I_{bkg}(q) \tag{3}$$

Assuming we've perfectly subtracted the background signal, the incident and reflected neutron beams are subject to Poisson noise. Therefore the variance in reflected and incident intensity is given by,

$$var(I_0) = I_0, \quad var(I_{refl}(q)) = I_{refl}(q) \tag{4}$$

Consequently, the error model of the reflectivity is given by the ratio distribution of two Poisson random variables. For large enough mean values, these Poisson distributions can be approximated by Gaussian. Under certain conditions,[1] the ratio distribution $Z = \frac{X}{Y}$ of two Gaussian distributed random variables $X$ and $Y$ , is again a Gaussian distribution with mean $\mu_z = \frac{\mu_x}{\mu_y}$ and variance, $\sigma_z$ given by,

$$\sigma_z^2 = \frac{\mu_x^2}{\mu_y^2}\left(\frac{\sigma_x^2}{\mu_x^2} + \frac{\sigma_y^2}{\mu_y^2}\right) \tag{5}$$

Using the variances for the incident and reflected beam from eqn. 4 in eqn. 5, we get, the variance in $R(q)$, as

$$var(R(q)) = \frac{R(q)}{I_0} + \frac{R(q)^2}{I_0} \tag{6}$$

To check the validity of eqn. 6 we compare it with the numerical results obtained using explicit Poisson distributed intensities, for the reflected and incident neutron beams, with mean and variance given by eqn.4. As shown in Figure S5 our noise model is in good agreement with the numerical results for $I_0 \geq 10^3$, which is well within the experimental noise range. Therefore for values typical to experiments, our noise model works well.

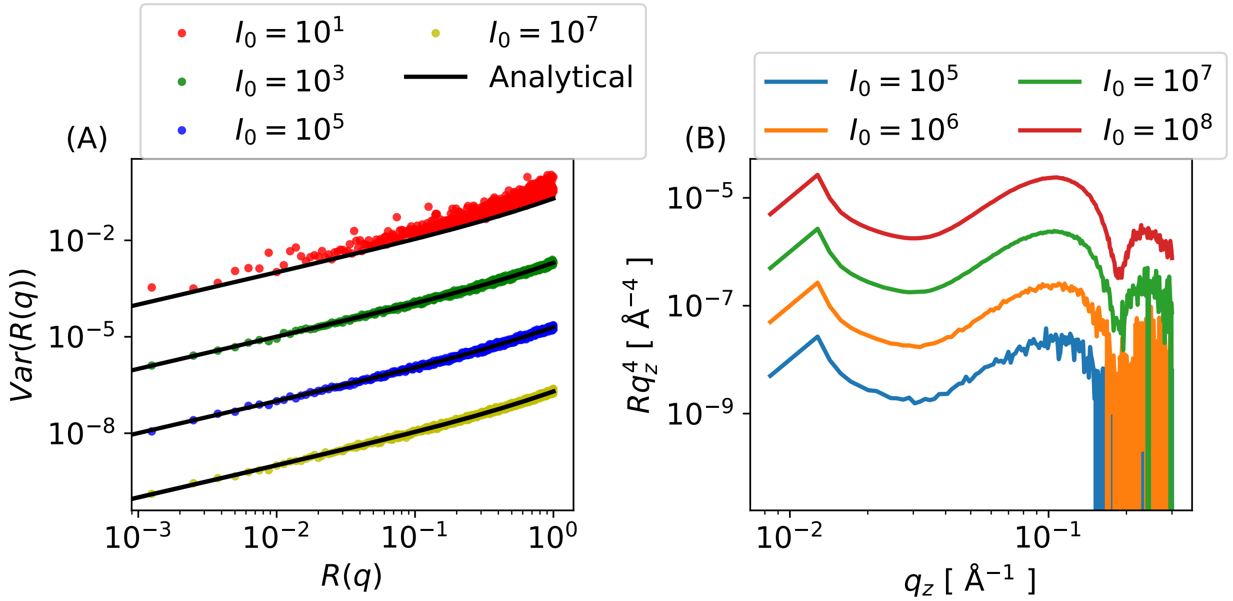


Figure S5: **Noise model for reflectivity.** (A) Comparison of analytical variances (Eqn. 6) with variances generated assuming Poisson distribution for different values of $I_0$ which controls the noise levels and can be considered as similar to incident intensity. (B) Neutron reflectivity profiles from a pure DOPC bilayer supported on silicon substrate with hLipid/$D_2O$ contrast at different noise levels. The reflectivity profiles at different $I_0$ values are arbitrarily shifted in $y-$direction for clarity.

### 3 Assigning Neutron Scattering Factors /Electron /Mass to CG-Martini Beads

Table S1: Example martini mapping file showing part of DOPC mapping file. The mapping file is taken from Ref.[2] (see also Martini mapping files in the lipidome.)

| **Index** | **Atoms** | **CG-Beads** |
|---|---|---|
| 1 | N | NC3 |
| 2 | C12 | NC3 NC3 NC3 PO4 |
| 3 | C13 | NC3 |
| 4 | C14 | NC3 |
| 5 | C15 | NC3 |
| 6 | H12A | NC3 NC3 NC3 PO4 |
| 7 | H12B | NC3 NC3 NC3 PO4 |
| .. | ... | ... |

Based on Table S1, $b_i$ for CG-bead $NC3$ is assigned as,

$$b_i(NC3) = b_i(N) + \frac{3b_i(C)}{4} + 3b_i(C) + \frac{3b_i(H)}{4} + \frac{3b_i(H)}{4} + ... \tag{7}$$

Where $b_i(X)$ is the neutron scattering factor/number of electrons or mass of atom $X$. We provide scripts for such assignment at https://git.rz.uni-augsburg.de/cbio-gitpub. The script takes as input the simulation `.gro` or `.pdb` file and path to the Martini mapping files.

In case of DOPC, the electron density (or mass density) obtained using above assignment method without backmapping leads to an artificial increase in the center of the profile (main text Figure 2). However, the neutron SLD does not exhibit such artifacts, therefore both direct and backmapping with energy minimization and 2 ps MD leads to the same SLD profile (Figure S6 A). This is most likely due to the negative scattering factor for hydrogen which leads to a fruitful cancellation of densities i.e higher density regions will also have more hydrogen.

For lipid with relatively shorter tails e.g DPPC the above mentioned artifact does not arise in the electron density profile i.e both the direct and backmapped method results in very similar profiles (Figure S6 B).

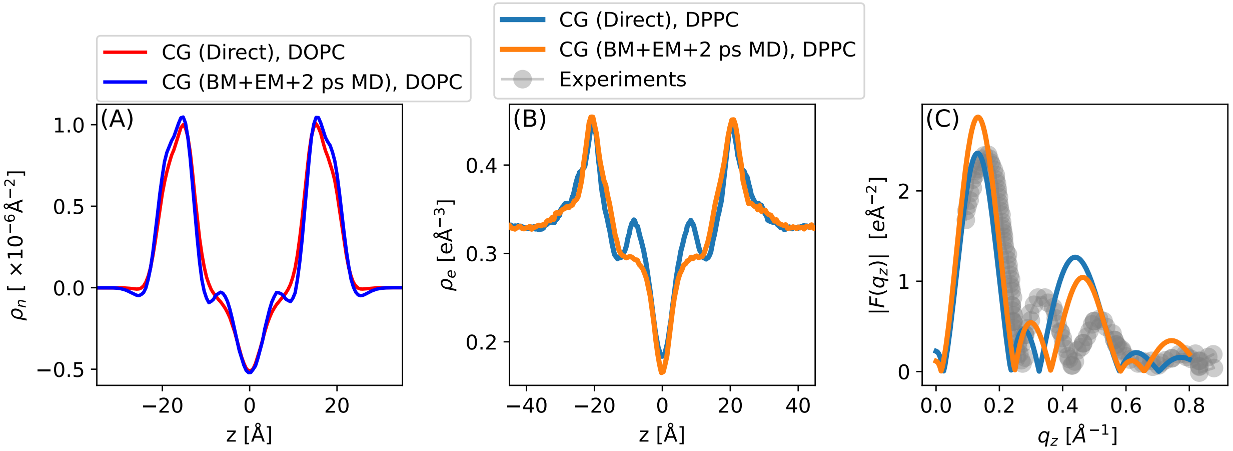


Figure S6: **Neutron SLD and electron density for CG-systems** (A) Neutron SLD of DOPC from direct CG method and from backmapping with energy minimization followed by a short 2 ps MD equilibration. (B) Electron density for DPPC from same two methods as in (A). (C) Corresponding X-ray scattering form factor for DPPC and comparison with experiments.[3] In this case (i.e DPPC), the agreement with experimental data is not as good as for DOPC (Figure S8 B).

### 4 Undulation Correction

The undulation correction is performed as described in Ref.[4] The implementation can be found at https://git.rz.uni-augsburg.de/cbio-gitpub.

1. For each frame consider a set of atoms e.g the phosphate or any carbon atom of lipid and obtain the Fourier coefficients as:

$$u(\mathbf{q}) = \frac{1}{2N} \sum_{k=1}^{N} \left[ z_{1k} e^{-i\mathbf{q}.\mathbf{r_{1k}}} + z_{2k} e^{-i\mathbf{q}.\mathbf{r_{2k}}} \right] \tag{8}$$

   $N$ is the number of lipid molecules per leaflet, $z_{1k}$ and $z_{2k}$ are the $z-$coordinates of each atom in leaflet 1 and leaflet 2 and $\mathbf{q} = 2\pi \left( \frac{\pm m}{L_x}, \frac{\pm n}{L_y} \right)$ with $L_x$ and $L_y$ being the box dimension in $x$ and $y$ directions.

2. Apply a filter of the type:

$$G(q) = \frac{1}{1 + (q/q_0)^4} \tag{9}$$

The coefficients become: $\tilde{u} = G(q/q_0)^{1/2}u$, such that the spectral intensity given by $\tilde{S} = S_u(q)G(q/q_0)$, where $S(u) = N\langle(|u(q)|^2)\rangle$.

The undulation spectra for the undulatory part should decay as $q^{-4}$ in accordance with predictions from continuum membrane models and hence the exponent 4 in eqn. 9. The higher $q-$modes corresponds to the detailed molecular structural motions and hence should be filtered out. The spectral intensity with and without the filter considering the terminal carbons (TC) and head group phosphorous atoms (P) is shown in Figure S7. As expected the lower $q$ modes follow $q^{-4}$.

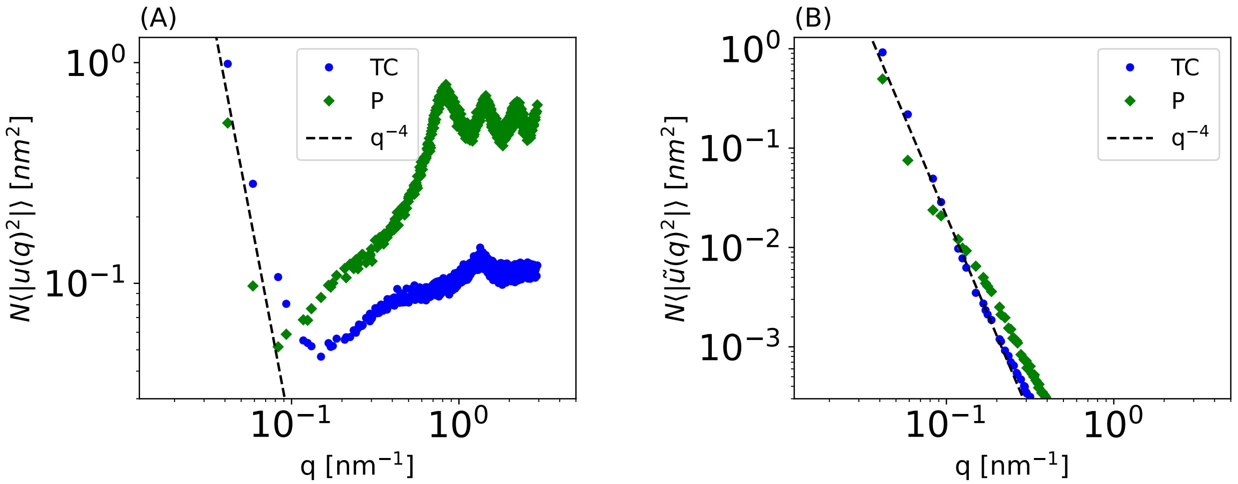


Figure S7: **Undulation spectra for DOPC bilayers** (A) Undulation spectra without applying filter. The low $q$ regime corresponding to undulation follow $q^{-4}$. (B) Undulation spectra after applying the filter (eqn. 9) to (A) using $q_0 = 0.08$ nm$^{-1}$. The calculations are done using either the terminal carbons (TC) shown in blue or the phosphorous atoms (P) shown in green. These plots agree well with the reported behavior.[4,5]

Electron density calculation via binning is now done with respect to the following surface,

$$z_k^{ref} = (z_k - \tilde{u}(r_k))\cos\theta \tag{10}$$

Where, $\tilde{u}(r_k) = \sum_q \tilde{u}(q)e^{i\mathbf{q}\cdot\mathbf{r}}$ is the inverse Fourier transform and $\theta$ denotes the local orientation of the plane and is obtained using the gradient of $\tilde{u}(r_k)$. The local orientation is obtained from $\tilde{\mathbf{n}} = (n_x, n_y, n_z) = \frac{(-\Delta\tilde{u},1)}{\sqrt{(\Delta\tilde{u})^2+1}}$, where, $\Delta\tilde{u}$ is the gradient of $\tilde{u}$. So, $\cos\theta = n_z(\mathbf{r_k})$

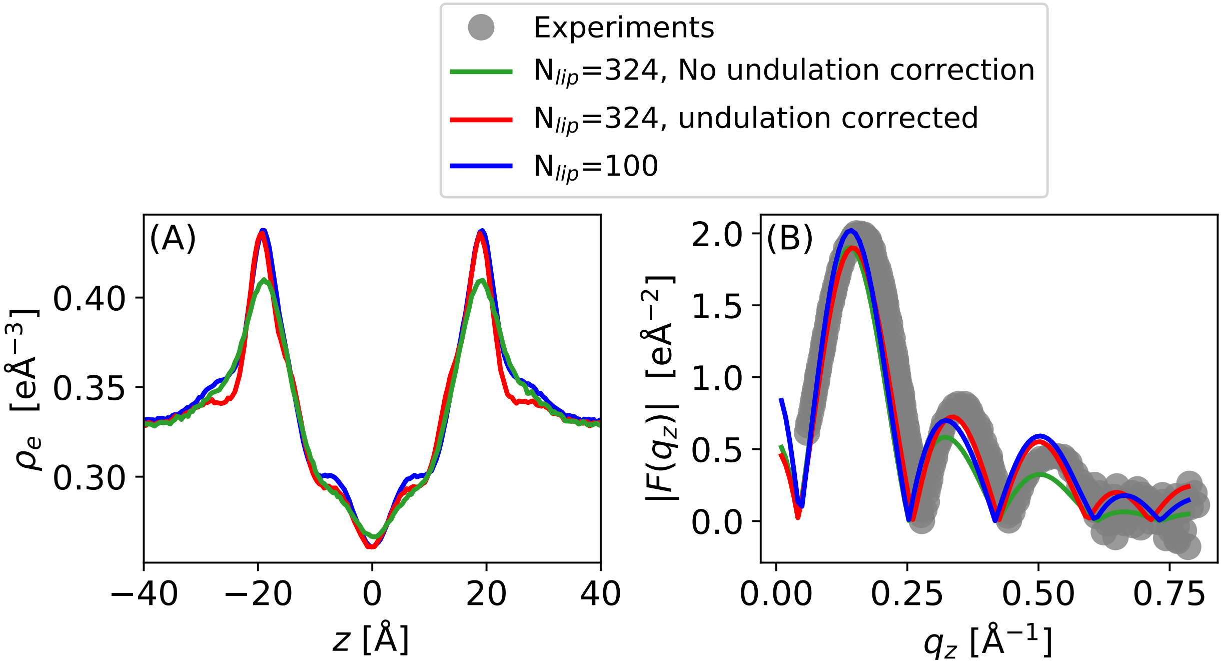


Figure S8: **Effect of bilayer undulations on the electron density.** (A) Electron density from bilayers of different sizes. For larger size (N=324 lipids/leaflet) the effect of undulation is depicted. (B) Corresponding X-ray form factors and comparison with experimental data from Ref.[3]

### 5 RNA Secondary Structure and Contact Frequency Maps

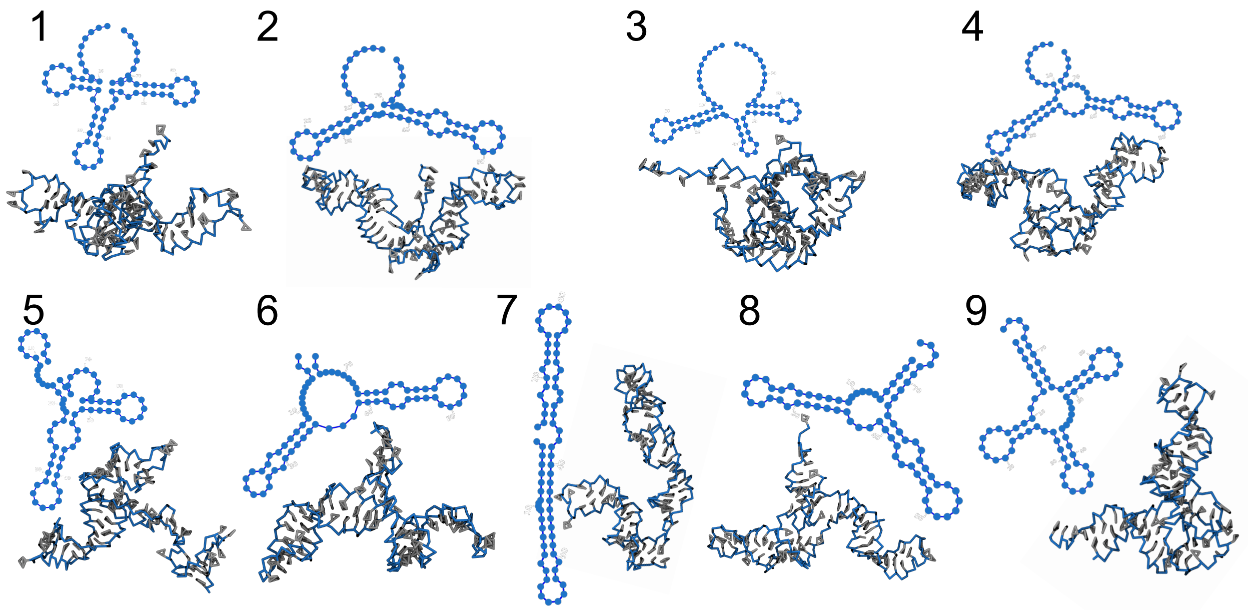


Figure S9: **tRNA secondary and corresponding tertiary structures**. Different secondary and corresponding tertiary structures of tRNA used in the simulations with neutral DOPC and cationic DOTAP bilayers. The secondary structures were predicted using the RNAfold (Vienna) package[6] and the corresponding tertiary structures were obtained using the RNAComposer webserver.[7]

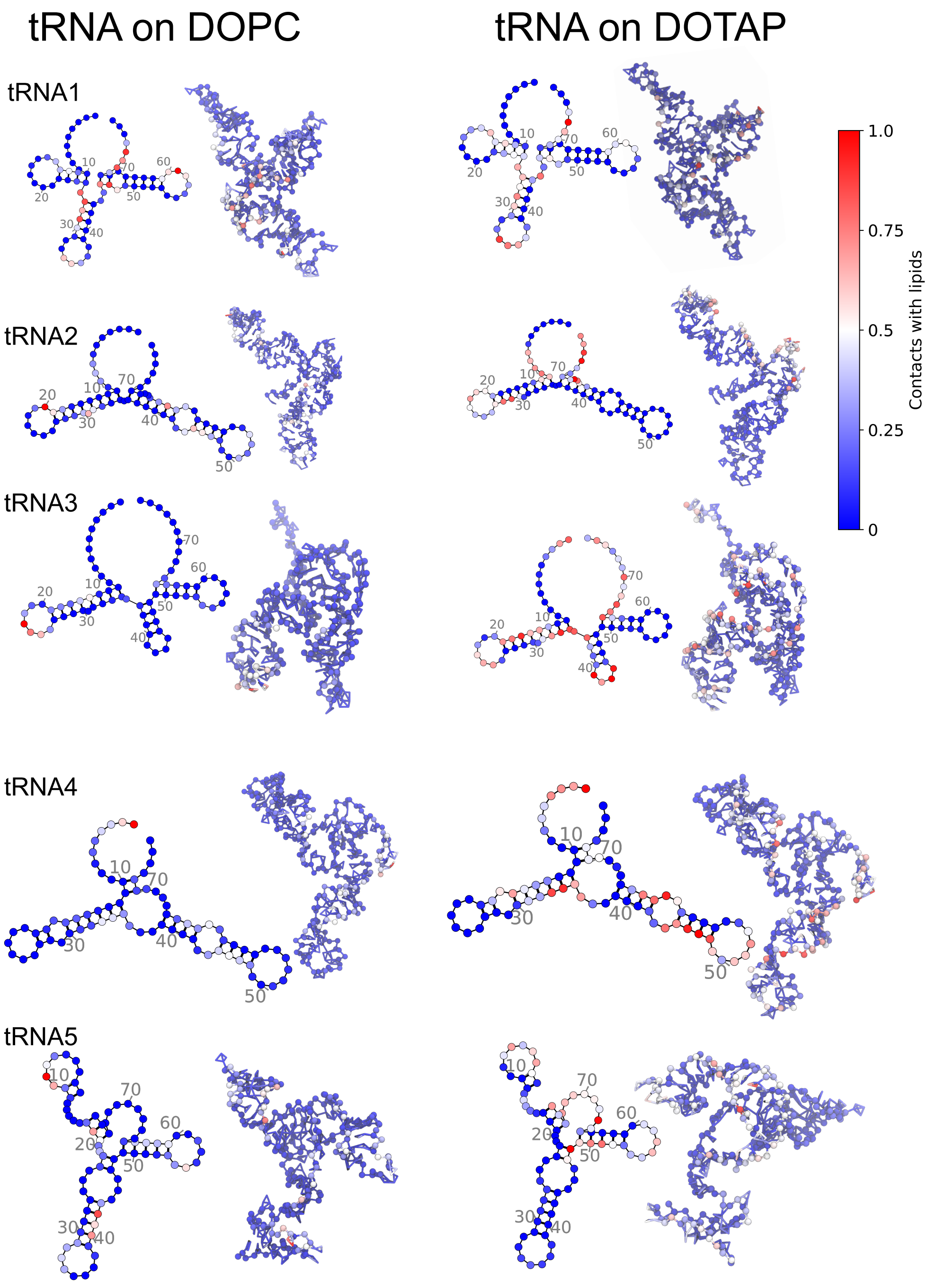


Figure S10: **RNA-Lipid contact frequency**. Contact frequency of tRNAlipid interactions on DOPC and DOTAP bilayers from CG simulations. For each bilayer type, the tRNA secondary structure and corresponding 3D conformation are shown. The contact frequencies are scaled relative to their respective maxima, so that the normalized values vary from 0 to 1.

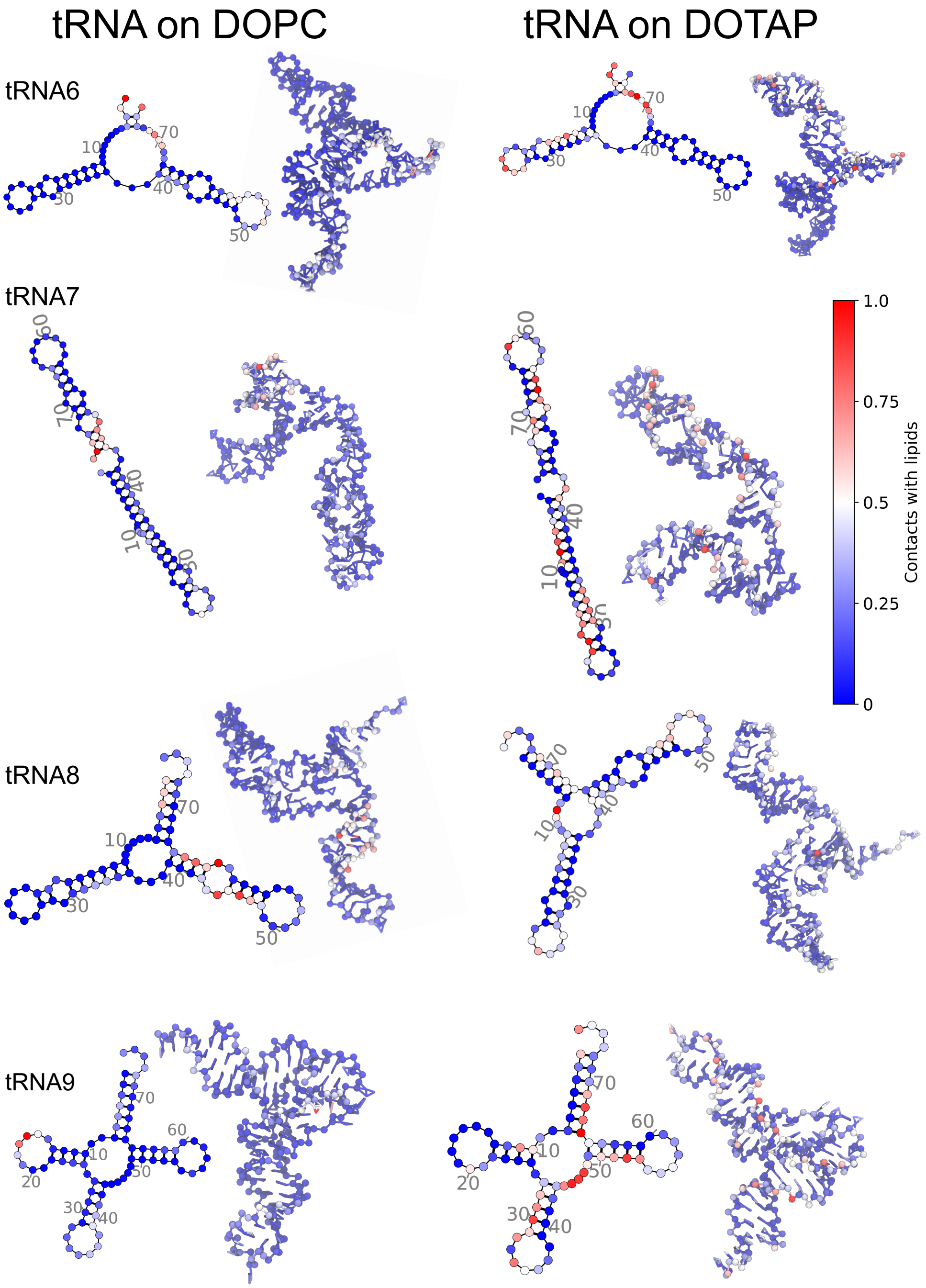


Figure S11: **RNA-Lipid contact frequency**. Same as in Figure S10 for the indicated tRNA conformations.

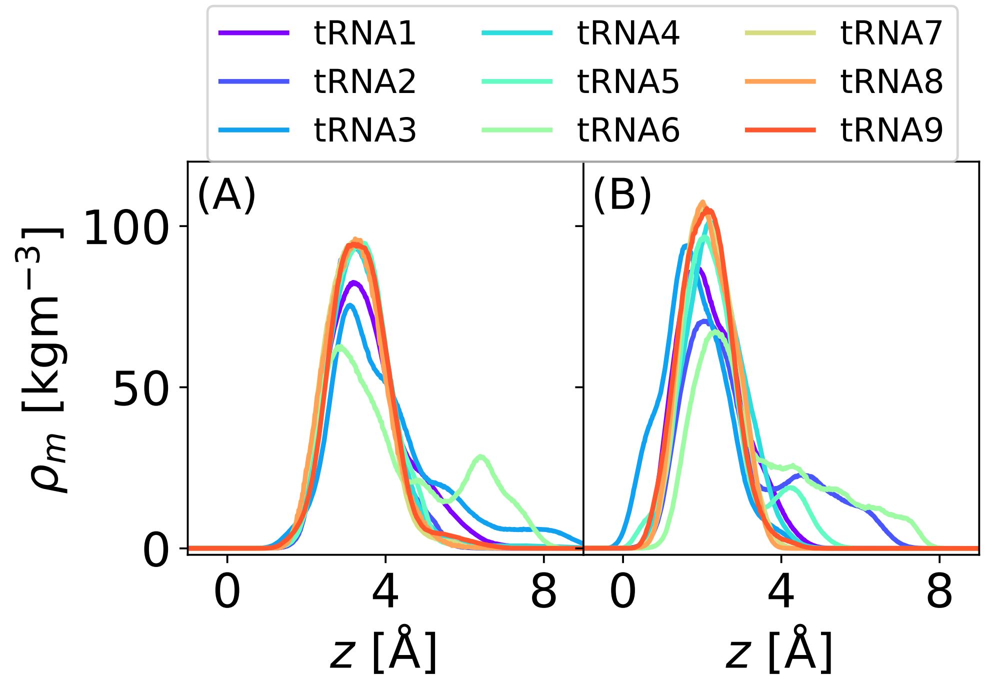


Figure S12: **Mass density profiles of different tRNA secondary structures along the bilayer normal.** The $z$ coordinate denotes the distance from the bilayer center. (A) Profiles for various tRNA secondary structures on DOPC bilayer and (B) corresponding profiles on DOTAP bilayer.

### 6 Additional Simulations for tRNA6 and tRNA9 on DOPC and DOTAP

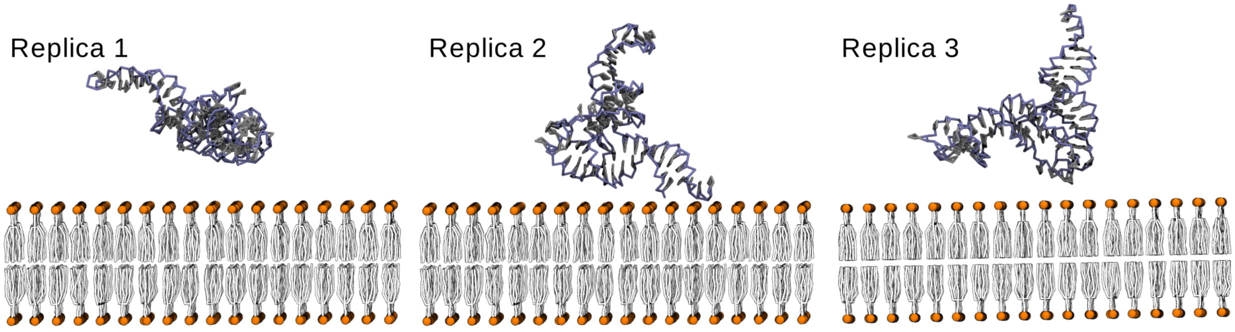


Figure S13: **Initial orientation of tRNA9 w.r.t the bilayer for different replicas.** In replica 1, the tRNA is oriented along the principal axis. Replica 2 and replica 3 are obtained by rotating the tRNA configuration in replica 1 by 90° about X-axis and Y-axis, respectively.

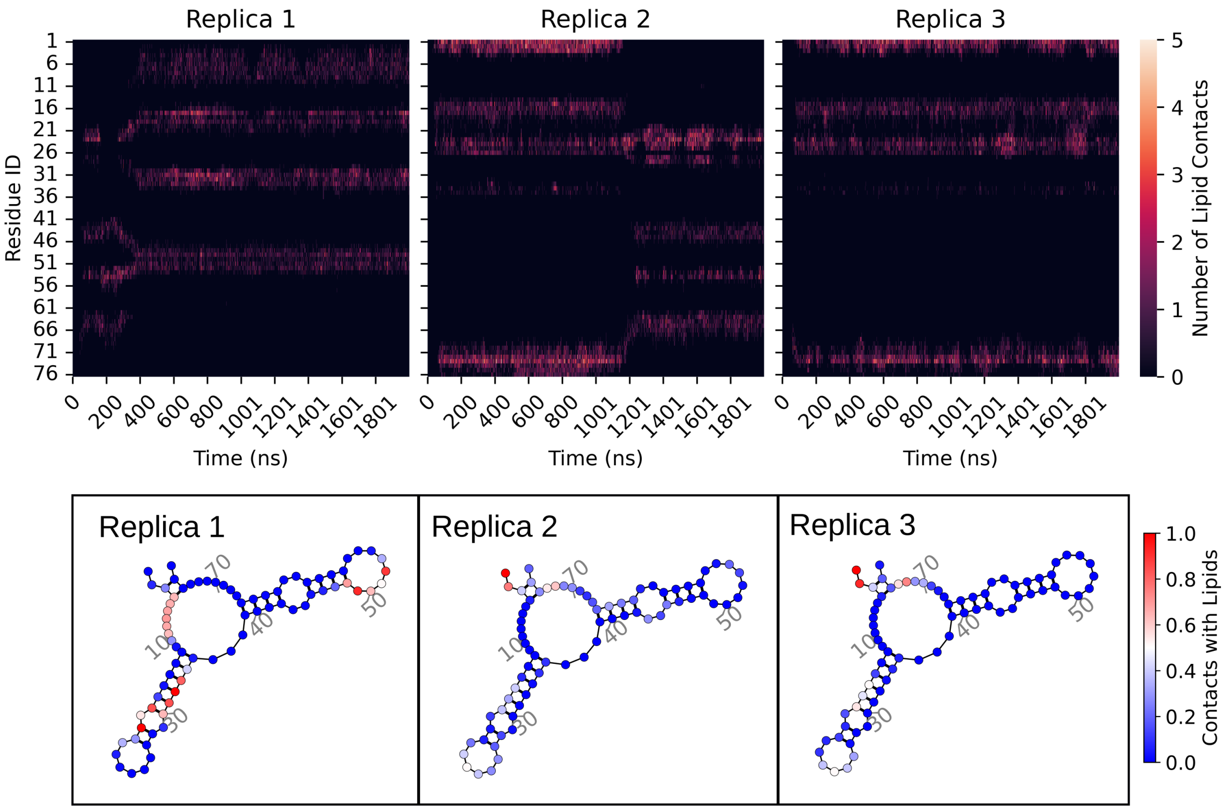


Figure S14: **Contact map for tRNA6 on DOPC.** Top row: Contact evolution of each tRNA residue with time over the whole simulation trajectory of 2 $\mu s$. The number of contacts for a tRNA residue at a given time is defined as the number of lipid headgroup beads (PO4 and NC3) within 5 Å of the residue. Bottom row: Number of contacts for each residue projected on the secondary structure. The number of contacts are normalized to the maximum observed value.

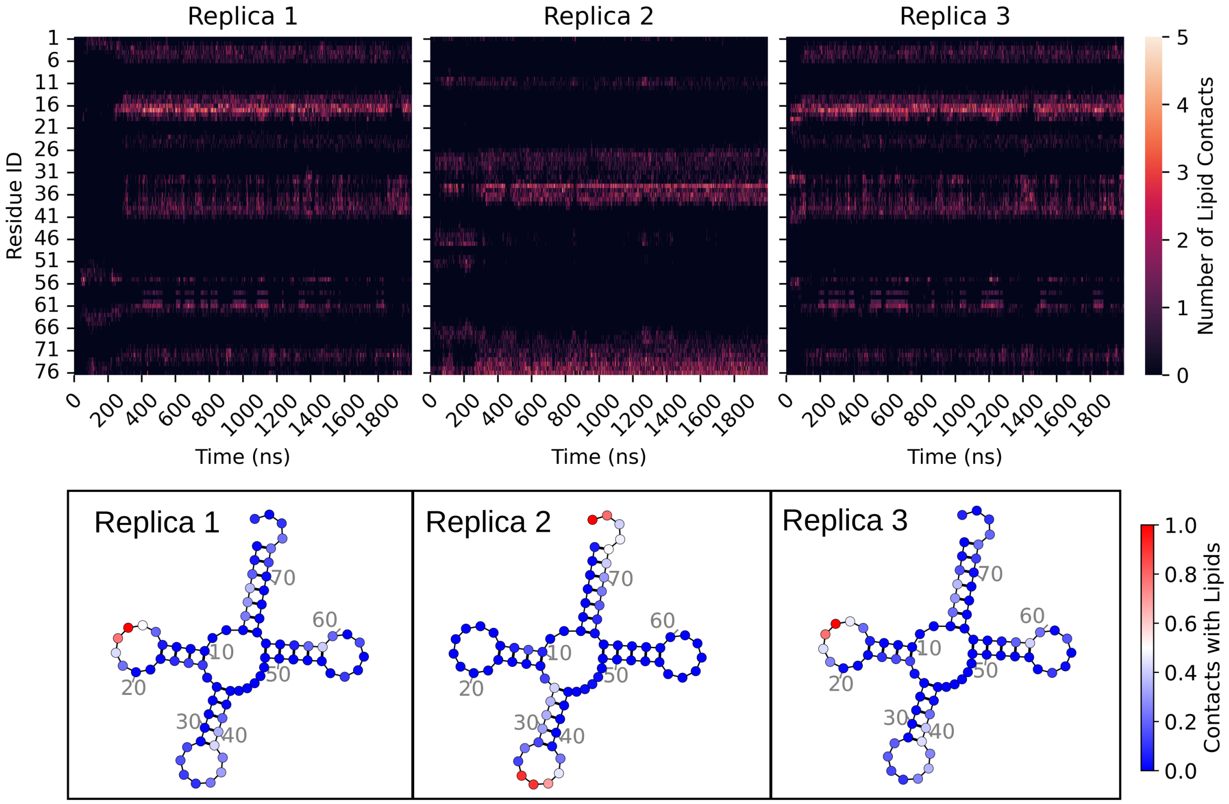


Figure S15: **Contact map for tRNA9 on DOPC.** Top row: Contact evolution of each tRNA residue with time over the whole simulation trajectory of 2 $\mu s$. The number of contacts for a tRNA residue at a given time is defined as the number of lipid headgroup beads (PO4 and NC3) within 5 Å of the residue. Bottom row: Number of contacts for each residue projected on the secondary structure. The number of contacts are normalized to the maximum observed value. Here, replica 1 and replica 3 gives rise to almost identical contact maps which is also evident from the top row.

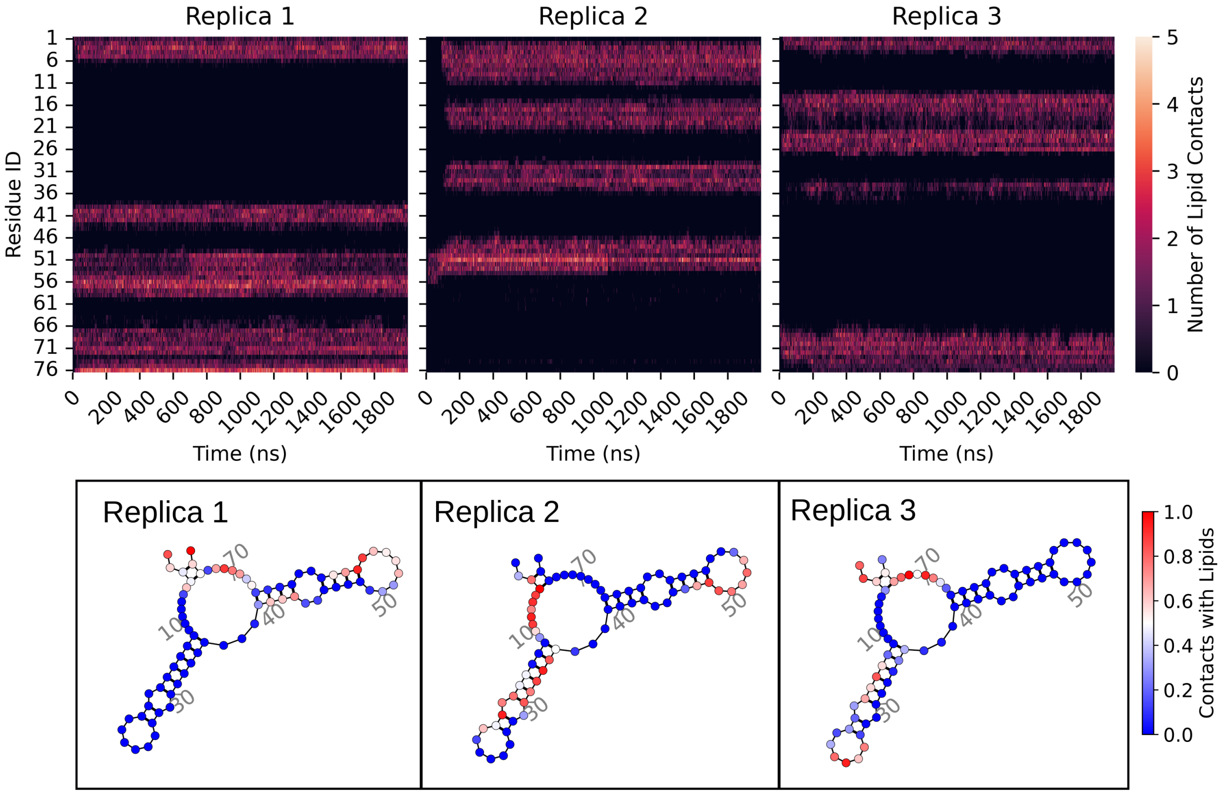


Figure S16: **Contact map for tRNA6 on DOTAP.** Top row: Contact evolution of each tRNA residue with time over the whole simulation trajectory of 2 $\mu s$. The number of contacts for a tRNA residue at a given time is defined as the number of lipid headgroup beads (PO4 and NC3) within 5 Å of the residue. Bottom row: Number of contacts for each residue projected on the secondary structure. The number of contacts are normalized to the maximum observed value.

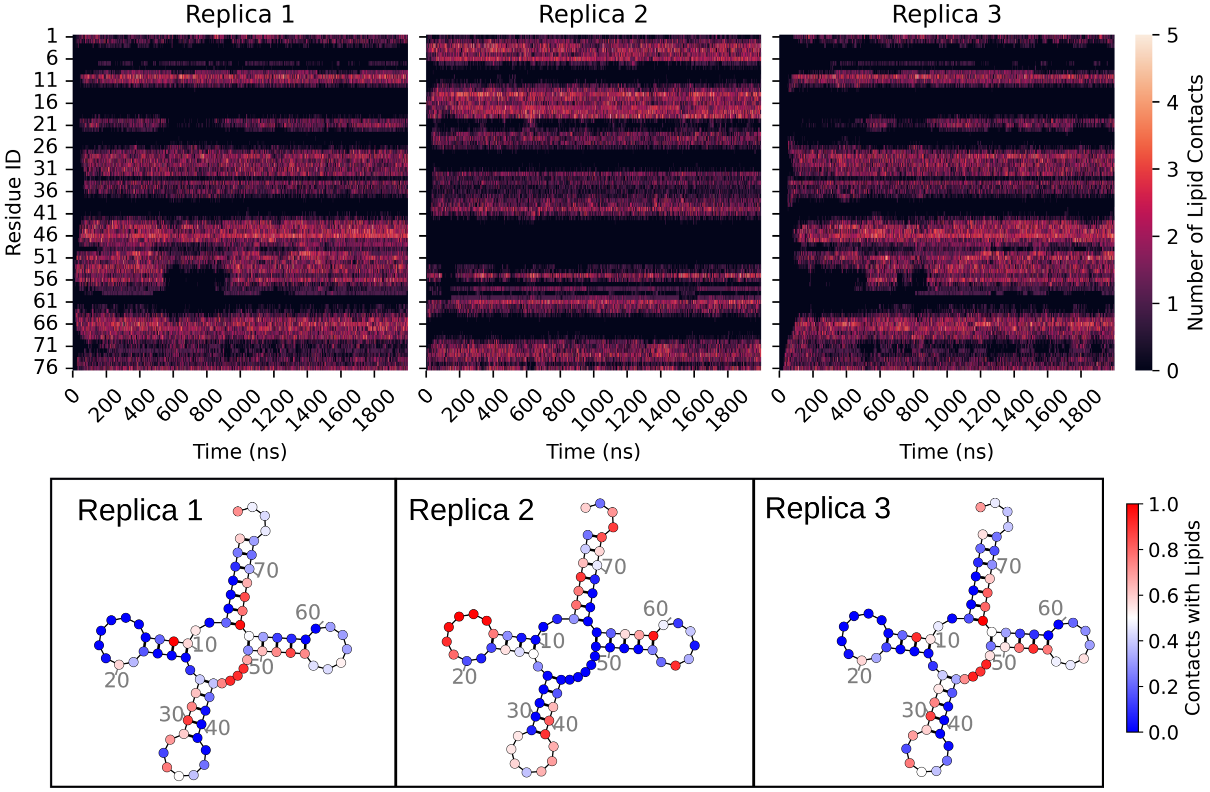


Figure S17: **Contact map for tRNA9 on DOTAP.** Top row: Contact evolution of each tRNA residue with time over the whole simulation trajectory of 2 $\mu s$. The number of contacts for a tRNA residue at a given time is defined as the number of lipid headgroup beads (PO4 and NC3) within 5 Å of the residue. Bottom row: Number of contacts for each residue projected on the secondary structure. The number of contacts are normalized to the maximum observed value.

### 7 NR Curves for tRNA at Bilayers and Monolayers

#### 7.1 tRNA/DOPC Systems

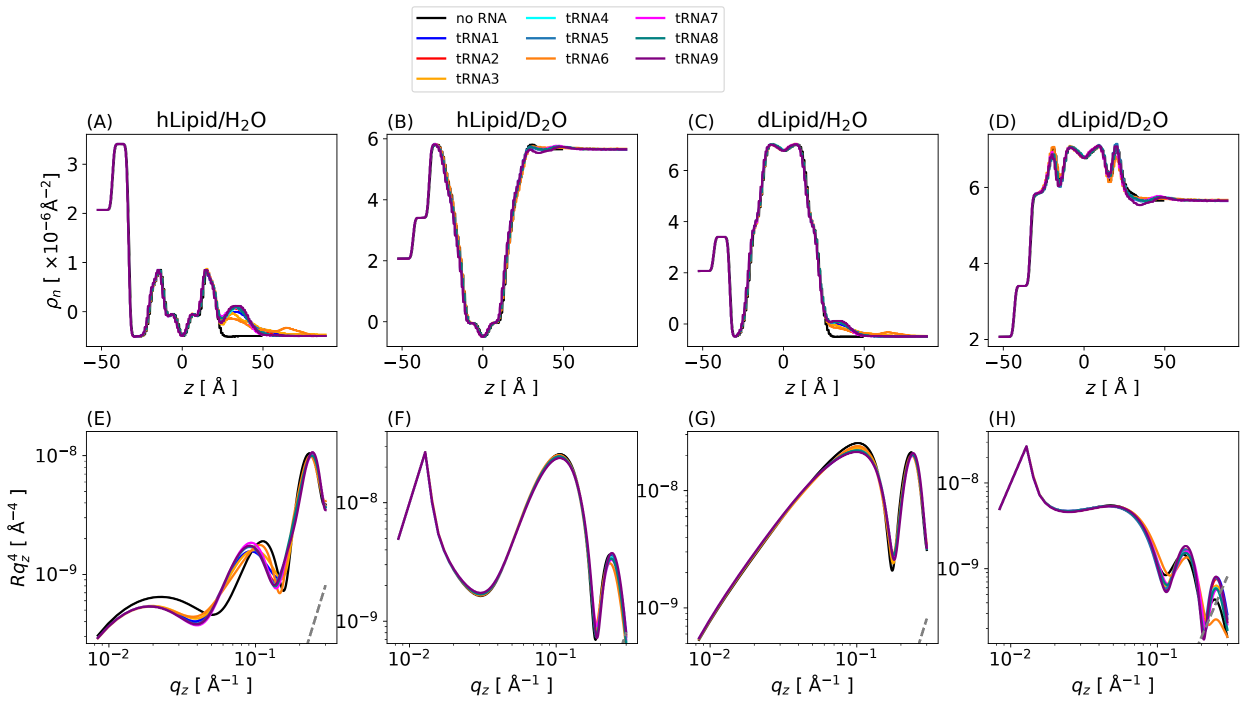


Figure S18: **Neutron SLD and NR profiles using four different deuteration conditions at the DOPC bilayer.** SLD for (A) non-deuterated solvent ($H_2O$) with regular non-deuterated lipid (hLipid), (B) deuterated solvent ($D_2O$) with hLipid, (C) $H_2O$ with deuterated lipid (dLipid), and (D) $D_2O$ with dLipid. (E-H) Corresponding NR profiles. The gray dashed line indicates the background signal at ($10^{-7}q_z^4$). Here, $L/N = 1.5$ in all cases.

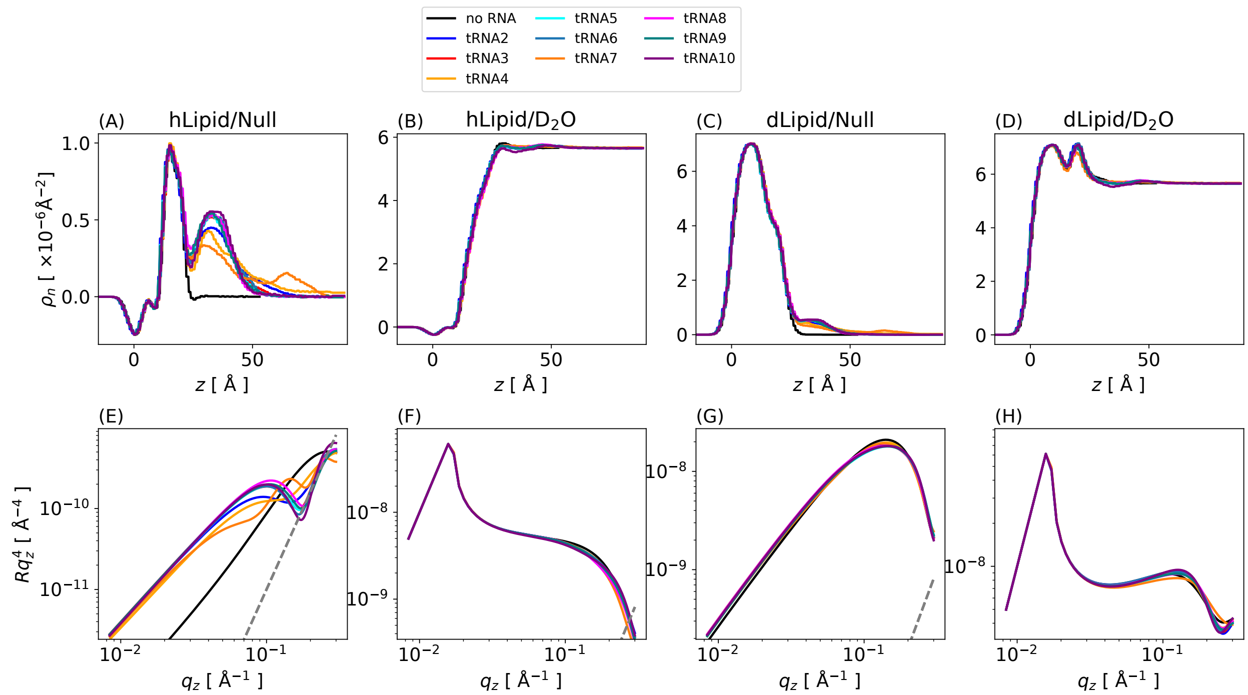


Figure S19: **Neutron SLD and NR profiles using four different deuteration conditions at the DOPC monolayer.** SLD for (A) non-deuterated solvent ($H_2O$) with regular non-deuterated lipid (hLipid), (B) deuterated solvent ($D_2O$) with hLipid, (C) $H_2O$ with deuterated lipid (dLipid), and (D) $D_2O$ with dLipid. (E-H) Corresponding NR profiles. The gray dashed line indicates the background signal at ($10^{-7}q_z^4$). Here, $L/N = 1.5$ in all cases.

#### 7.2 tRNA/DOTAP Systems

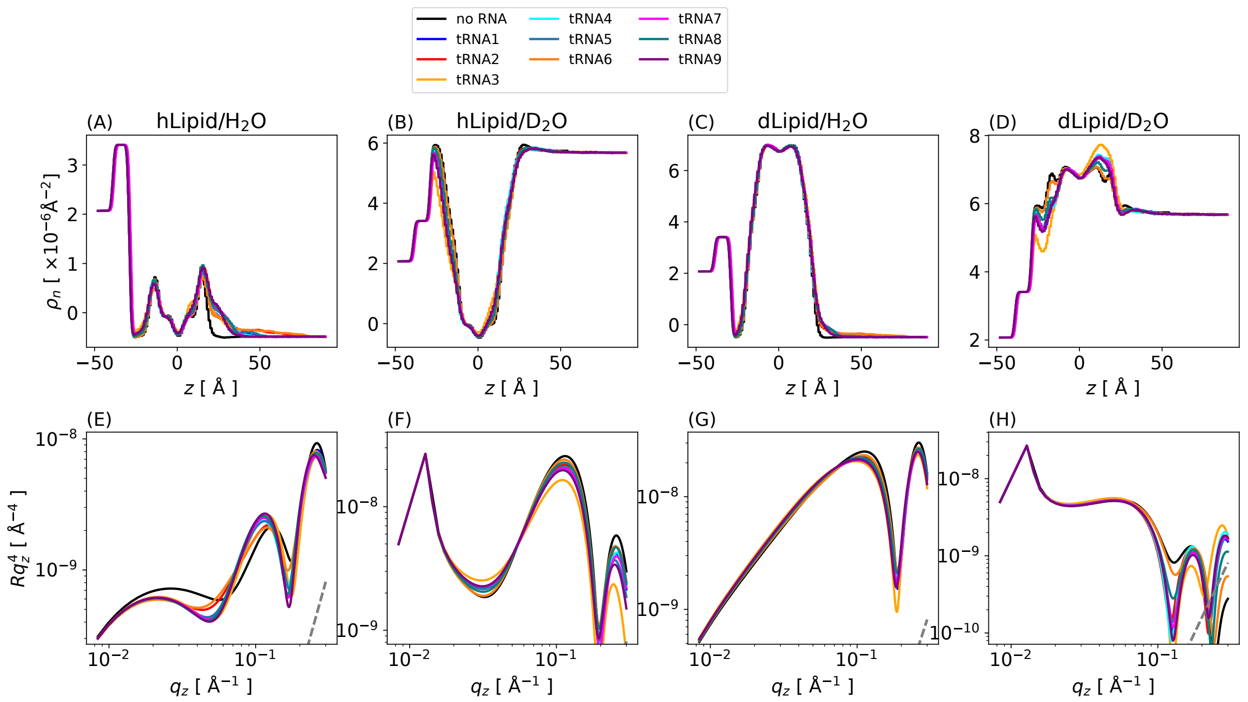


Figure S20: **Neutron SLD and NR profiles using four different deuteration conditions at the DOTAP bilayer.** Conditions as in Figure S11.

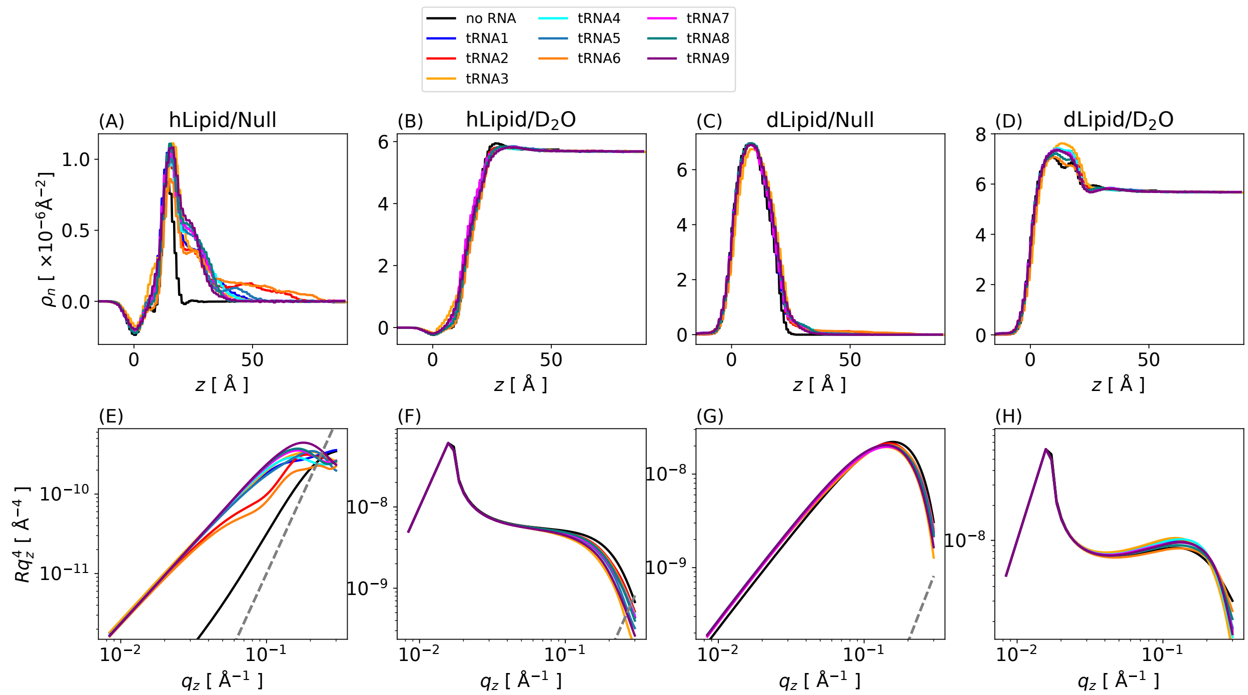


Figure S21: **Neutron SLD and NR profiles using four different deuteration conditions at the DOTAP monolayer.** Conditions as in Figure S12.

### 8 XRR for RNA at Bilayers and Monolayers

#### 8.1 tRNA/DOPC Systems

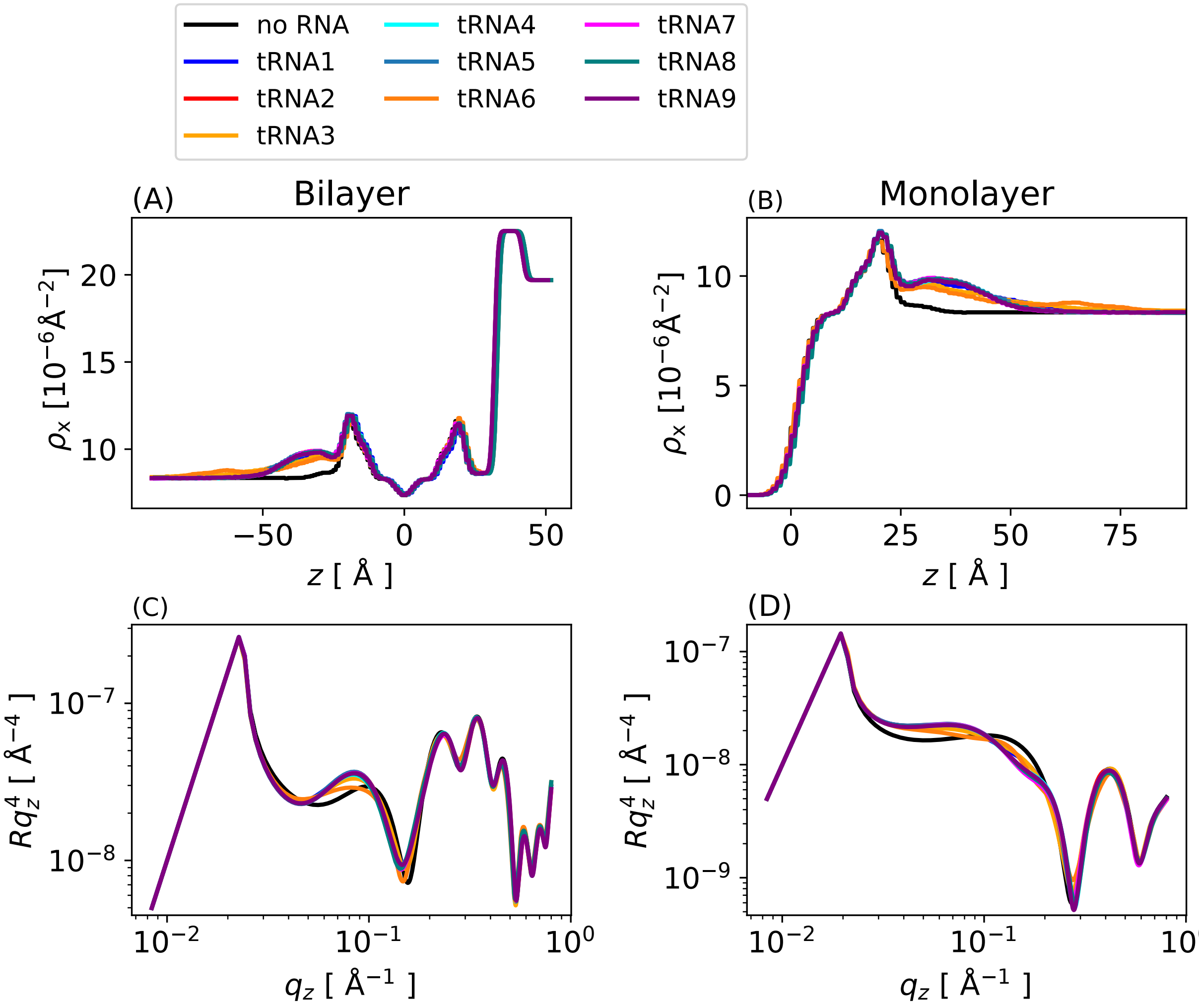


Figure S22: **X-ray SLD and XRR profiles for DOPC bilayers (A, C) and monolayers (C, D)**. Results are for $L/N = 1.5$ and for all the tRNA secondary structures. The line depicting the background noise level is omitted since its much lower than the signal.

#### 8.2 tRNA/DOTAP Systems

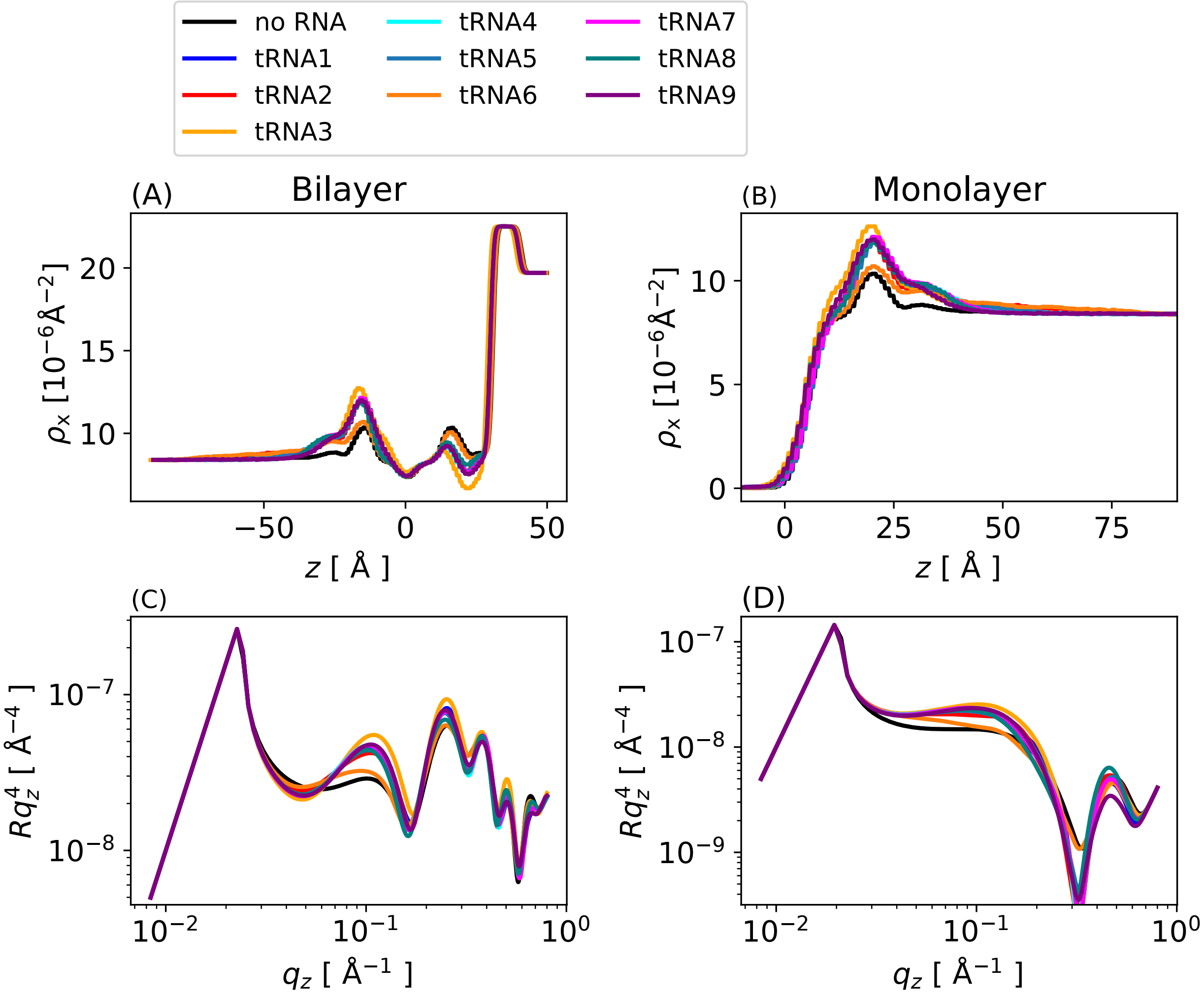


Figure S23: **X-ray SLD and XRR profiles for DOTAP bilayers (A, C) and monolayers (C, D)**. Same conditions as in Figure S15.

### 9 Lipid-to-Nucleotide Ratio Dependence of NR and XRR Signals

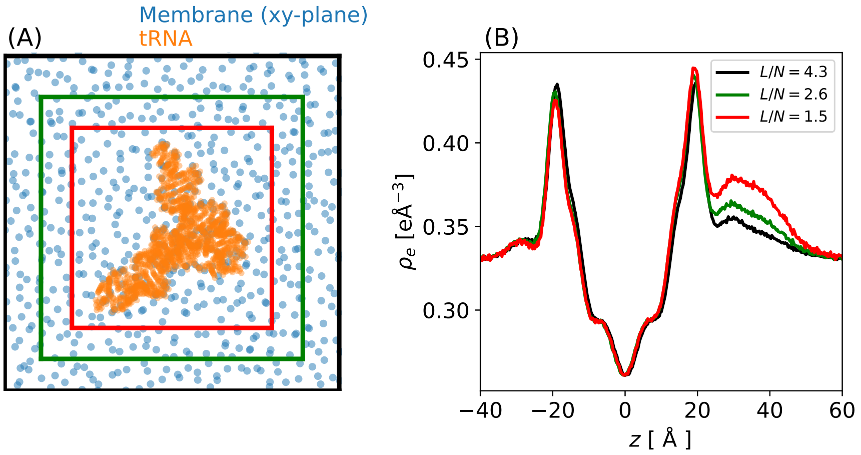


Figure S24: **Variation of the lipid-to-nucleotide ratio.** (A) To include the effect of RNA concentration, we assume different areas around the tRNA for the SLD calculation. Here the rectangle around RNA corresponds to $L/N = 4.2$ (black), $L/N = 2.6$ (green), and $L/N = 1.5$ (red). The corresponding electron density is shown in (B). Lowering $L/N$ increases the contribution of RNA to the electron density between 25 Å $< d <$ 50 Å.

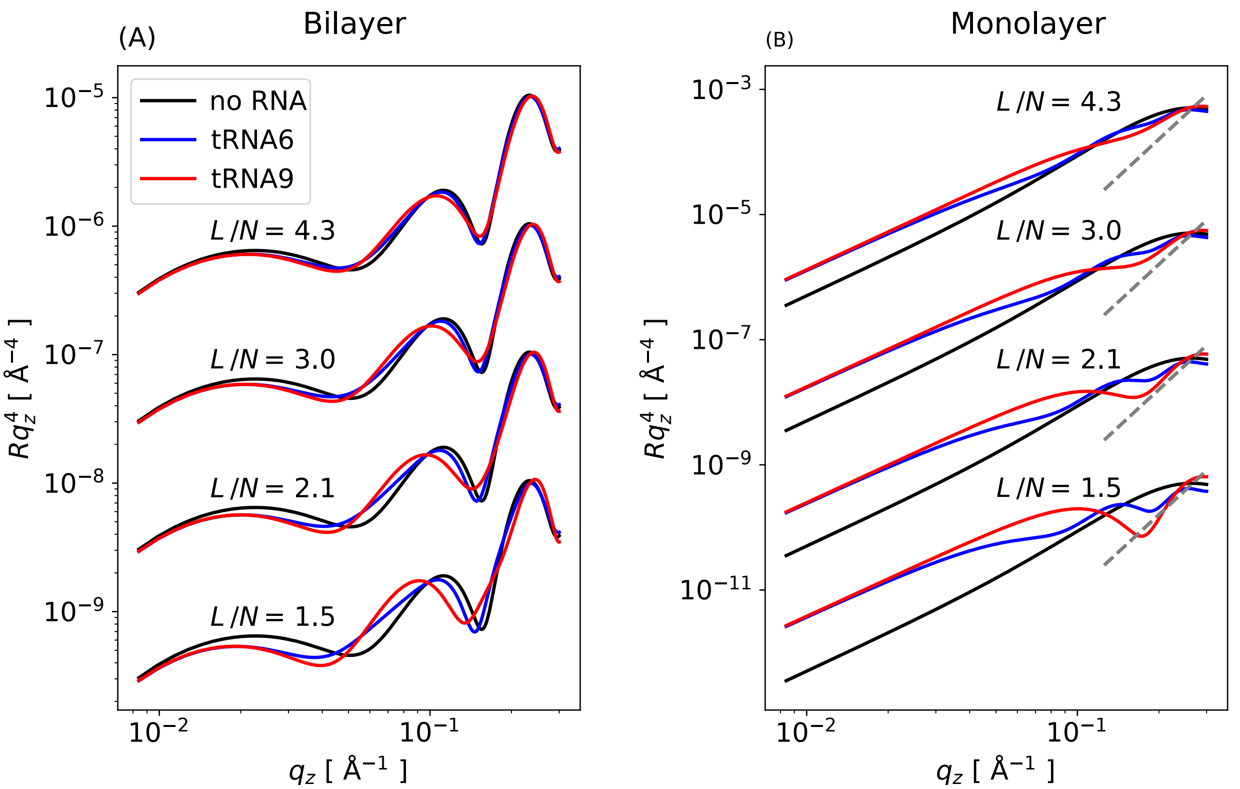


Figure S25: **Influence of $L/N$ ratio on NR profiles.** (A) NR for DOPC bilayer and monolayers (B). The profiles for different $L/N$ values are shifted by arbitrary constant along the $y-$axis. The gray dashed line indicates the background signal at ( $10^{-7}q_z^4$). For (A) the background is much lower than the signal and hence omitted.

.

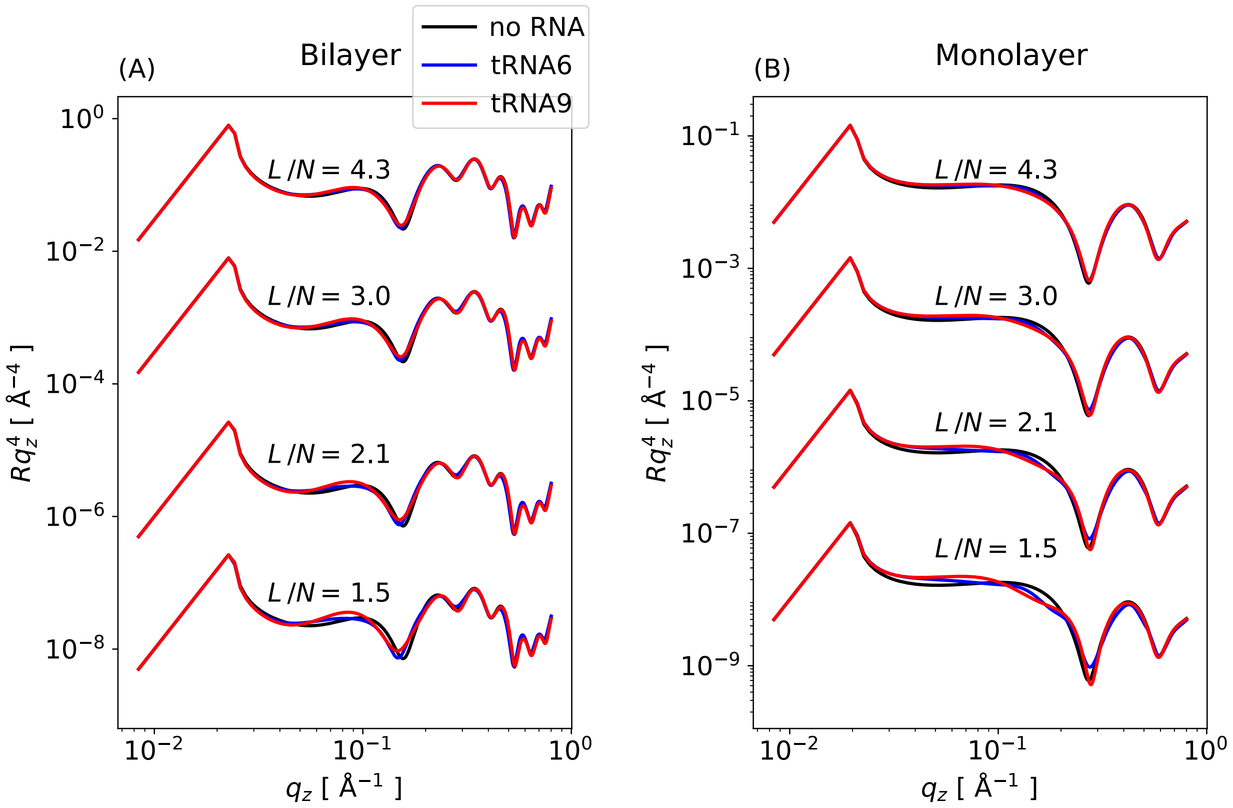


Figure S26: **Influence of $L/N$ ratio on XRR profiles.** (A) XRR for DOPC bilayer and monolayers (B). The profiles for different $L/N$ values are shifted by arbitrary constant along the $y-$axis. The line depicting the background noise level is omitted here since it is much lower than the signal.

.
